# Ultraviolet attenuates centromere-mediated meiotic genome stability and alters gametophytic ploidy consistency in flowering plants

**DOI:** 10.1101/2024.02.12.579936

**Authors:** Huiqi Fu, Jiaqi Zhong, Jiayi Zhao, Li Huo, Chong Wang, Dexuan Ma, Wenjing Pan, Limin Sun, Ziming Ren, Tianyi Fan, Ze Wang, Wenyi Wang, Xiaoning Lei, Guanghui Yu, Jing Li, Yan Zhu, Danny Geelen, Bing Liu

## Abstract

Ultraviolet (UV) radiation influences development and genome stability in organisms; however, its impacts on meiosis, a special cell division essential for the delivery of genetic information over generations in eukaryotes, remain not yet elucidated. In this study, we demonstrate that UV attenuates the centromere-mediated meiotic chromosome stability and induces unreduced gametes in *Arabidopsis thaliana*. We show that UV reduces crossover (CO) rate but does not interfere with meiotic chromosome integrity. Functional centromere-specific histone 3 (CENH*3*) is required for the obligate CO formation, and plays a role in protection of homolog synapsis and sister-chromatid cohesion under UV stress. Moreover, UV specifically alters the orientation and organization of spindles and phragmoplasts at meiosis II, resulting in meiotic restitution and unreduced gametes. Further, we determine that UV-induced meiotic restitution does not rely on the UV Resistance Locus8-mediated UV perception and the Tapetal Development and Function1- and Aborted Microspores-dependent tapetum development, but occurs possibly via impacted JASON function and downregulated Parallel Spindle1. This study sheds light on the impacts of UV on meiotic genome stability and gametophytic ploidy consistency, which thus may influence genome evolution in flowering plants.

## Introduction

Meiosis is a specialized type of cell division, which involves one round of chromosomal DNA replication and twice chromosome segregation, leading to production of gametes with a halved ploidy. At prophase I, meiotic recombination takes place between homologous chromosomes, which is initiated by the formation of DNA double-strand breaks (DSB) (Hartung et al., 2007). Meiotic DSBs are repaired by RAD51-mediated intersister-chromatid recombination and/or the DMC1-catalyzed homologous recombination (Da Ines et al., 2013; Kurzbauer et al., 2012). Defects in DSB repair result in failed crossovers (COs) between homologs and production of univalents, and/or chromosome fragmentation (Sanchez-Moran et al., 2007; Su et al., 2017). During meiotic recombination, homologous chromosomes undergo pairing and synapsis, which involves assembly of the synaptonemal complex (SC) between the juxtaposed homologs (Gordon and Rog, 2023). The lateral and central elements of SC are composed of the HORMA domain protein HOP1/ASY1 and the transverse filament protein ZYP1, respectively (Armstrong et al., 2002; Higgins et al., 2005). SC assembly is required for the obligate CO formation (France et al., 2021), which drives reciprocal exchange of genetic materials and creates genetic diversity, and is required for the faithful segregation of homologous chromosomes (Wang and Copenhaver, 2018; Zickler and Kleckner, 2023).

Programmed chromosome segregation is controlled by dynamic assembly and organization of microtubule- and actin-composed cytoskeleton (Roeles and Tsiavaliaris, 2019; Sofroni et al., 2020; Xu et al., 2013). At metaphase I, chromosomes are aligned at the cell plate by a bipolar spindle, which pulls the homologous chromosomes to the opposite poles at anaphase I; at metaphase II, two spindles are formed and drive the separation of sister chromatids at anaphase II. Lesions in the orientation or formation of spindle and/or phragmoplast may lead to irregular nuclei positioning and defective cytokinesis with resultant formation of unreduced gametes, or cause unbalanced chromosome separation and aneuploids (Andreuzza et al., 2015; Brownfield et al., 2015; De Storme and Geelen, 2011; Niu et al., 2015).

A key chromosome element required for the maintenance of chromosome stability and normal dynamics is the centromeres, where the multi-complex structure kinetochore is patched on and physically links the chromosomes to microtubules (Verdaasdonk and Bloom, 2011). In addition to playing a canonical role as the sites of spindle attachment, centromeres facilitate multiple meiosis processes including the protection of the centromeric cohesion and sister kinetochore co-orientation in meiosis I (Brar and Amon, 2008). In plants, the centromeres safeguard chromosome stability at both normal and high temperatures, and this trait has been used for haploid induction (Ahmadli et al., 2023; Jin et al., 2023; Peterson et al., 2010; Ravi and Chan, 2010; Wang et al., 2021b). Nevertheless, the role of the centromeres in meiosis at stressful environmental conditions awaits further studies.

It has been evidenced that whole genome duplication (WGD) broadly occurs in higher plants, which drives speciation, diversification, and adaption to environment (Ren et al., 2018; Soltis et al., 2015; Van de Peer et al., 2020). Formation and fusing of unreduced gametes derived from non-reductional meiotic cell division, called meiotic restitution, are considered the main route to WGD (Ramsey and Schemske, 1998). Meiotic restitution can be triggered by either genetic alterations or temperature stresses, which cause loss of meiosis cell cycle, aberrant organization of spindle and/or phragmoplast or irregular cytokinesis (De Storme et al., 2012; De Storme and Geelen, 2011; De Storme and Geelen, 2020; Lei et al., 2020; Ravi et al., 2008; Zhou et al., 2022). However, whether meiotic restitution can be triggered by other environmental factors is less understood.

Ultraviolet is an invisible form of light, which has short wavelength and situates in the portion of the electromagnetic spectrum between visible light and X-rays (Diffey, 2002). Based on the wavelength, UV spectrum is further designated into UV-A (400–315 nm), the long-wave UV that can pass through the atmosphere and is considered less toxic because it is not absorbed by native DNA; UV-B (315–280 nm), which carries more energy than UV-A and is responsible for the radiation’s best-known effects on organisms; and UV-C (280–100 nm), the high-energy UV that is entirely screened out by the stratospheric ozone and thus does not reach Earth’s surface (Diffey, 2002). Exposure to UV radiation is unavoidable for plants when capturing sunlight for photosynthesis. UV affects many development processes in plants, such as endoreduplication, hypocotyl and cotyledon development, flavonoid accumulation, and DNA stability (De et al., 2022; Li et al., 2013; Ries et al., 2000; Shi and Liu, 2021; Verdaguer et al., 2017). In addition, UV influences plant development by inducing reactive oxygen species (ROS) and antioxidants (Gill et al., 2015; Sinha and Häder, 2002).

UV induces multiple DNA lesions including cyclobutane-pyrimidine dimers (CPDs) and 6-4 photoproducts (6-4PPs), and their Dewar valence isomers and DNA strand breaks (Garinis et al., 2005; Gill et al., 2015; Hessling et al., 2021; Mullenders, 2018; Rastogi et al., 2010; Schalk et al., 2017; Sinha and Häder, 2002; Takahashi et al., 2011). In mammals, it has been evidenced that CPDs and 6-4PPs need to be properly repaired through the conserved nucleotide excision repair (NER) pathway, otherwise DSBs can be induced via the blocked and consequent abortion of DNA replication (Garinis et al., 2005; Oh et al., 2011; Thoma, 1999; Wakasugi et al., 2014). However, the impact of UV on the genome stability in sexual reproductive cells has not been elucidated.

It is a common feature that the reproductive development in plants, especially gametogenesis, is sensitive to UV (Çetinbaş-Genç et al., 2022; Feng et al., 2000; Koti et al., 2004; Sampson and Cane, 1999; Torabinejad et al., 1998). During gametogenesis, the composition of sporopollenin, which composes the outer cell walls of pollen grains and microspores, is prone to be altered by UV radiation (Çetinbaş-Genç et al., 2022; Del Valle et al., 2020; Xue et al., 2020; Yang et al., 2023). The UV-sensitivity of pollen is widely used as a monitor of UV flux, or a recorder of the past environment changes, or for reading the signature of flower evolution (Bell et al., 2018; Benca et al., 2018; Fraser et al., 2014; Fraser et al., 2011; Willis et al., 2011; Zhang et al., 2014). To date, however, little is known how UV may affect meiosis.

Environmental pollution has been destructing the stratospheric ozone layer, which increases the possibility of extreme UV exposure and thus boosts the risk of diseases such as cancer and the threat to agriculture safety (Barnes et al., 2019; Benca et al., 2018; Hessling et al., 2021). It thus is important to address how UV may affect reproduction and genome stability in plants. In this study, we demonstrate that UV interferes with centromere-mediated chromosome stability and induces meiotically-restituted unreduced gametes in Arabidopsis. Our findings unveil the influences of UV on the meiotic genome stability and gametophytic ploidy stability, which may subsequently affect genome evolution in higher plants.

## Results

### UV radiation induces unreduced gametes formation in *Arabidopsis thaliana*

To explore the impact of UV on male reproduction in plants, we treated flowering *Arabidopsis thaliana* wild-type Columbia (Col) with UV-C for 12 or 8 h, and pollen grain formation was examined using Alexander staining. Flower buds at 1- and 2-day post treatment (1 and 2 dpt) did not show an obvious difference in viable pollen formation compared with control (Supplemental Fig. S1). However, locules in the anthers of flowers at 3 and 4 dpt showed an obvious reduction in viable pollen grain formation, but no viable pollen grain was yielded in flowers at 5 dpt (Fig. 1B; Supplemental Fig. S1). In addition, treatment by UV-C for as short as 1 h induced about 50% aborted pollen grains (Fig. 1B). These data indicated that UV-C radiation disrupts gametogenesis. Interestingly, enlarged pollen grains, yet inviable revealed by FDA staining, were occasionally observed in flowers at 5-7 dpt (lower than 0.10%) (Fig. 1A and C). We suspected that the impaired gametogenesis by UV-C may lower the rate of the enlarged pollen grains; hence, we tried UV-B or UV-A radiation, which have longer wave and lower energy than UV-C, for 24 h. We found that the UV-B- and UV-A-radiated Col at 7 dpt exhibited low levels of abortion of pollen development (Fig. 1A and B) and generated 4.74% and 6.79% enlarged pollen grains, respectively, which can germinate on pollen medium and thus were viable (Fig. 1A, C and E).

**Figure 1.**
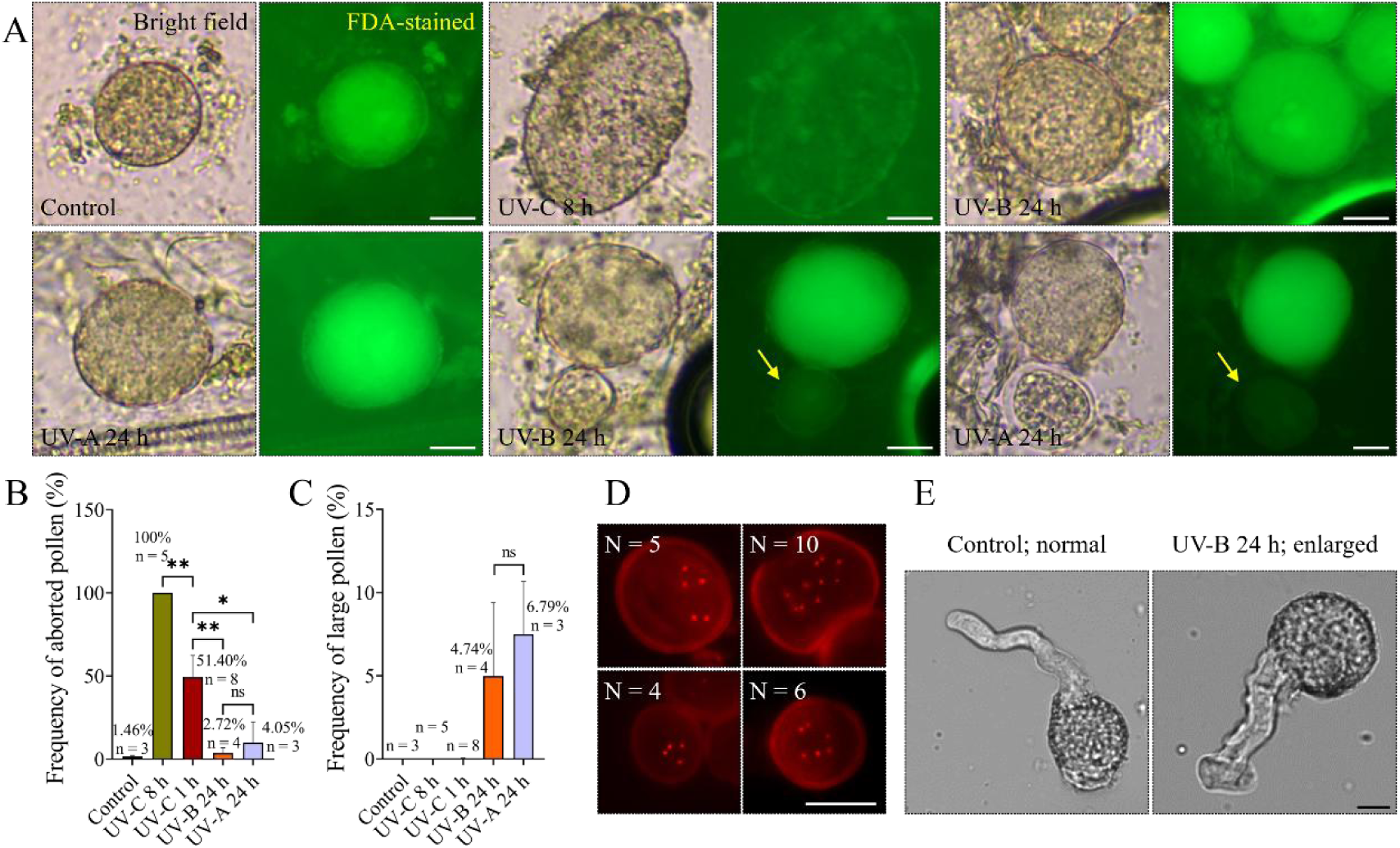
UV induces unreduced gametes in *Arabidopsis thaliana*. A, Pollen grains produced by Col at control condition, or radiated by UV. Yellow arrows indicate inviable pollen grains. B, Graph showing the frequencies of aborted pollen grains in UV-radiated Col at 5 dpt. C, Graph showing the frequencies of larger pollen grains in UV-radiated Col at 7 dpt. Unpaired *t*-tests were performed; frequencies indicate the rates of the corresponding phenotypes; n indicates the numbers of analyzed plant individuals; ** indicates *P* < 0.01; * indicates *P* < 0.05; ns indicates *P* > 0.05. D, Representative live-imaging of unicellular-staged microspores in Col expressing *pCENH3::TagRFP-CENH3* at control condition, or radiated by UV. N indicates the numbers of RFP-CENH3 foci. E, Germination of pollen grains with a normal size from control plants, and enlarged pollen grains yielded by UV-B-radiated Col. Scale bars = 10 μm.

The enlarged size of pollen grains suggests an increased DNA content or ploidy (De Storme et al., 2013). To determine the exact chromosome number in the larger pollen grains, we took advantage of the *pCENH3::RFP-CENH3* reporter, which enables labeling and quantification of the centromeres (Komaki et al., 2020). At unicellular stage, a haploid microspore had five RFP-CENH3 foci, representing five chromosomes; in comparison, the plants radiated by either UV spectrum yielded microspores with ten RFP-CENH3 foci, indicating that they were diploid (Fig. 1D). Moreover, microspores with four or six RFP-CENH3 foci, which indicate aneuploids, were occasionally observed (Fig. 1D). Taken together, these data demonstrate that UV radiation induces unreduced gametes formation in Arabidopsis.

### UV evokes meiotic restitution and primarily induces FDR-type unreduced gametes

The doubled chromosome number in microspores at unicellular stage suggested that the ploidy diploidization occurred prior to gametogenesis, likely during meiosis (Fig. 1D). To verify this, we analyzed tetrad-staged meiocytes in Col upon UV radiation. A normal tetrad contained four similarly-sized spores with each harboring one haploid nucleus (Fig. 2A). In Col radiated by either UV spectrum for 8 h, meiotic products with irregular configurations were observed (Fig. 2A and B). These aberrant meiocytes included triads that contained one diploid spore and two haploid spores, dyads that contained two diploid spores, unbalanced dyads that contained one haploid spore and one triploid spore, and monads that contained four haploid spores, all of which manifested occurrence of meiotic restitution. In addition, unbalanced triads and tetrads, and polyads were also observed, yet at very low levels (Fig. 2A and B). The rates of the UV-B- and UV-A-induced meiotically-restituted meiocytes were largely elevated when the treatment was prolonged to 24 h (Fig. 2B; 58.33%, UV-B; 64.25%, UV-A). Analysis of *quartet* (*qrt*), in which four meiotic products are released as a tetrad configuration (Francis et al., 2006), confirmed formation of meiotically-restituted or aneuploid microspores upon UV radiation (Fig. 2C and D). In addition, Col radiated by shorter periods of UV-C also generated dyads and triads and unreduced microspores, which indicated a quick response of male meiosis to UV radiation (Supplemental Fig. S2). Moreover, both three UV spectrum induced microspores with rough cell walls (Fig. 2E), supporting that gametogenesis was perturbed by the UV radiation. Overall, these findings demonstrate that UV radiation induces male meiotic restitution.

**Figure 2.**
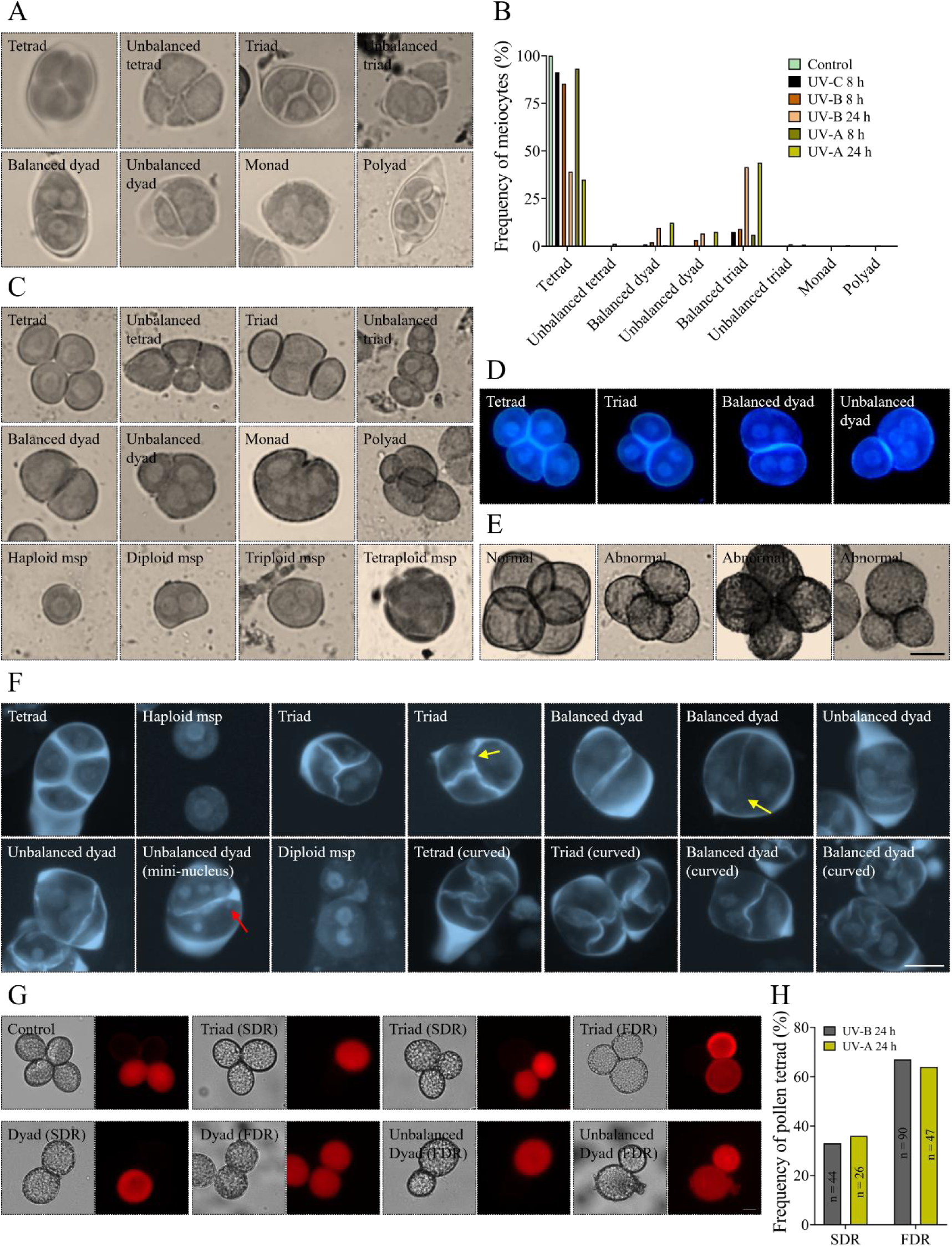
UV radiation primarily induces FDR-type meiotic restitution. A, Representative images of tetrad-staged meiocytes in Col with different configurations. B, Graph showing the rates of tetrad-staged meiocytes in Col at control condition or radiated by UV. C, Representative images of unicellular stage microspores in *qrt* with different configurations. D, DAPI-stained unicellular stage microspores in *qrt* with different configurations. E, Representative images of bi/tricellular stage microspores in *qrt* with normal or abnormal cell walls. F, Representative images of DAPI- and aniline blue-stained meiocytes at tetrad stage and unicellular stage microspores with different configurations in Col at control condition or radiated by UV. Red arrows indicate mini-nucleus; yellow arrows indicate incomplete callose cell walls. G, Representative images of different types of pollen tetrads in I3bc expressing RFP. H, Graph showing the rates of SDR- and FDR-type pollen tetrads induced by UV-B or UV-A. n indicates the number of the analyzed meiotically-restituted pollen tetrads. Scale bars = 10 μm.

A combination of aniline blue and 4’,6-diamidino-2-phenylindole (DAPI) staining was applied to check the positioning of nuclei and cell wall formation in the UV-induced aberrant meiotic products. Normal tetrads harbored four haploid nuclei, which were separated by callosic cell walls, and each spore developed into a haploid microspore (Fig. 2F). In Col radiated by either UV spectrum, we observed triads and dyads with either a complete loss of one or more cell walls and associated adjacent localization of nuclei and resultant diploid microspores, or breaks in the callosic cell walls (yellow arrows) (Fig. 2F). In line with the orcein staining, mini-nucleus were visualized (red arrow) (Fig. 2F). Moreover, some meiocytes showed curved callosic cell walls (Fig. 2F). These data indicate that meiotic cytokinesis is interfered by UV radiation.

To determine if the UV-induced unreduced gametes are derived from the first or second division restitution (FDR or SDR), we took advantage of the fluorescent-tagged lines (FTLs) in the *qrt* background, which carry T-DNA inserts encoding fluorescent proteins in mature pollen grains (Francis et al., 2007). Arabidopsis harboring a heterozygous status of the fluorescent reporter produces pollen tetrads, in which two pollen grains show fluorescence and two pollen grains do not (Fig. 2G). In a meiotically-restituted dyad or triad, if fluorescence appears both in a diploid spore and a haploid spore, or two diploid spores, or only in a triploid spore, or both in a haploid spore and a triploid spore, it suggests that homologous chromosomes are not separated and thus indicate FDR; in contrast, if fluorescence only appears in a diploid spore or two haploid spores, it indicates that sister chromatids do not disjoin, thus manifesting SDR (Fig. 2G).

Since UV-C impairs gametogenesis, we only evaluated meiotic restitution in plants radiated by UV-B or UV-A. The FTL reporter line I3bc was used for UV treatment, and the expression of RFP in pollen tetrads from fully-opened flowers at 6 or 7 dpt was examined. We found that both UV-B and UV-A primarily induced FDR, and the rates occurred at very similar levels (Fig. 2H; UV-B, 67.16%; UV-A, 64.38%). However, it should be noted that meiotic recombination events at the chromosome regions that cover the loci of the reporters may interfere with the segregation ratio; the frequencies we detected thus only reflect a proximity.

### UV induces aberrant nuclei positioning at meiosis II and increases chromosome instability in *cenh3*

To explore the cytological basis of the UV-induced meiotic alterations, we monitored meiotic chromosome behaviors in UV-radiated Col. In control plants, meiocytes at metaphase I showed five aligned bivalents, and at metaphase II, homologous chromosomes were distributed at two cell polars (Fig. 3A and B). At telophase II, four haploid chromosome sets were formed with each developing into haploid nucleus at tetrad stage (Fig. 3C and D). In UV-B- and UV-A-radiated Col, metaphase I meiocytes harboring univalents were occasionally observed, despite they did not occur at a significant level (Fig. 3E and Q). In UV-C-radiated plants, about 9.09% metaphase II meiocytes showed abnormal separation of sister chromatids (Fig. 3F and R), which, however, did not occur in Col under UV-B or UV-A (Fig. 3R). Notably, adjacent localization of isolated haploid chromosome sets or nuclei, which revealed a triad configuration, was frequently visualized at telophase II or tetrad stage in Col radiated by either UV spectrum (Fig. 3G and H). The UV-induced meiotic restitution thus was probably caused by the irregular nuclei positioning.

**Figure 3.**
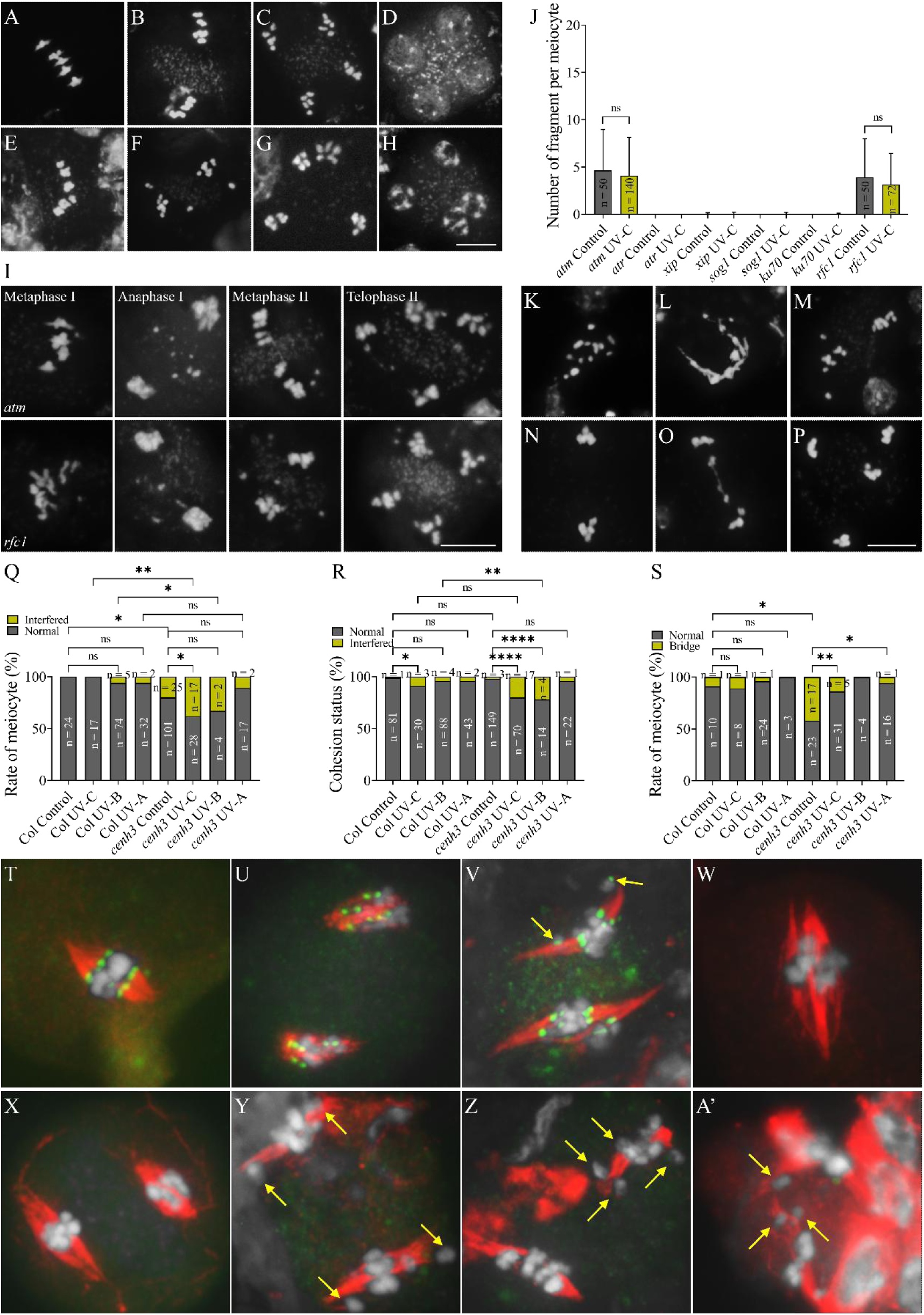
UV induces irregular nuclei positioning at meiosis II and increases meiotic instability in *cenh3*. A-H, DAPI-staining of chromosomes at metaphase I and II, telophase II and tetrad in Col at control condition (A-D) and radiated by UV (E-H). I, Representative images of DAPI-stained metaphase I, anaphase I, metaphase II and telophase II chromosomes in *atm* and *rfc1* at control condition or radiated by UV-C. J, Graph showing the average number of chromosome fragment per meiocyte in *atm*, *atr*, *xip*, *sog1*, *ku70* and *rfc1* at control and UV-C conditions. K-P, Representative images of DAPI-stained metaphase I (K and L), metaphase II (M), anaphase I (N and O) and telophase II (P) chromosomes in *cenh3* at control condition or radiated by UV-C. Q-S, Graphs showing the rates of meiocytes showing univalent formation at metaphase I (Q), interfered sister-chromatids segregation from metaphase I to II (R), or chromosome bridges at anaphase I (S) in Col and *cenh3* at control and UV conditions. T-A’, Co-immunolocalization of ɑ-tubulin and CENH3 in Col (T-V) and *cenh3* (W-A’) at control (T and U, W and X) and UV-C (V, Y-A’) conditions. White, DAPI; red, ɑ-tubulin; green, CENH3; yellow arrows indicate separated sister chromatids. Scale bars = 10 μm. For J, unpaired *t* tests were performed; for Q-S, χ^2^-tests were performed; n indicates the number of the analyzed cells; **** indicates *P* < 0.0001; ** indicates *P* < 0.01; * indicates *P* < 0.05; ns indicates *P* > 0.05.

The primary DNA lesions induced by UV-C are cyclobutane-pyrimidine dimers (CPDs) and 6-4 photoproducts (6-4PPs), which, if not repaired successfully through the conserved nucleotide excision repair (NER) pathway, can induce DNA double-strand breaks (DSBs) (Garinis et al., 2005; Oh et al., 2011; Thoma, 1999; Wakasugi et al., 2014). To address if UV-C induces DSB formation in Arabidopsis meiosis, we performed immunolocalization of ɤH2A.X protein, the phosphorylated form of H2A.X that specifically locates near DSB sites and thus is often taken as a DSB marker (Burma et al., 2001), and the DSB repair protein DMC1 on zygotene and/or pachytene chromosomes. We found that the foci formation of these two DSB marker proteins did not show an obvious alteration upon UV-C radiation (Supplemental Fig. S3A and B), hinting that UV-C may not induce ectopic DSB formation in Arabidopsis meiocytes. To verify this, we examined the impact of UV-C on meiotic chromosome stability in meiocytes of a series of DNA repair mutants, in which the mutated genes act in different DNA damage response pathways. First of all, Arabidopsis with dysfunction of the eukaryote-conserved kinase Ataxia Telangiectasia-Mutated (ATM), which plays a crucial role in DSB repair and participates in regulation of multiple meiosis processes (Paull, 2015), was analyzed. At normal condition, *atm* showed an average of 5.00 chromosome fragments per meiocyte (Fig. 3I and J), which indicate a defect in DSB repair as reported (Kurzbauer et al., 2021; Zhao et al., 2023). UV-C radiation for 8 h did not altered the number of chromosome fragments in *atm* (Fig. 3J). Arabidopsis with a loss-of-function of the ATM ortholog protein Ataxia Telangiectasia and Rad3-related (ATR) did not show a perturbed chromosome integrity at either control or UV-C condition (Fig. 3J). These data suggested that UV-C does not interfere with the ATM/ATR-dependent meiotic genome stability in Arabidopsis meiocytes. In support of this, dysfunction of either ɤH2A.X-Interacting Protein (XIP) or Suppressor of Gamma Response 1 (SOG1), which mediates DSB response via the RAD51-dependent recombination pathway and acts downstream of ATM/ATR to regulate various DNA damage responses, respectively (Fan et al., 2022; Li et al., 2022; Wei et al., 2021; Yoshiyama et al., 2017), showed normal chromosome integrity at both conditions (Fig. 3J). Likewise, *ku70*, which is hypersensitive to DSB-inducing agents due to attenuated nonhomologous end joining (NHEJ) pathway for DSB repair (Tamura et al., 2002), showed normal chromosome integrity (Fig. 3J). Moreover, we analyzed meiotic chromosome integrity in Arabidopsis with dysfunction of the conserved DNA replication protein Replication Factor C (RFC1), which plays a crucial role in meiotic DSB repair, and, in mammals, is required for the nucleotide excision repair pathway-mediated DNA damage response under UV radiation (Liu et al., 2010; Liu et al., 2013; Shiomi et al., 2012; Yu et al., 2012). At control condition, *rfc1* yielded univalents and chromosome fragments that manifest compromised crossover (CO) formation and DSB repair, respectively (Fig. 3I and J). The UV-C radiation, however, did not alter the level of chromosome fragmentation in *rfc1* (Fig. 3J). Taken together, these findings demonstrate that UV-C does not interfere with meiotic chromosome integrity in Arabidopsis.

In Arabidopsis, the maintenance of genome stability under high temperature conditions requires normal assembly and function of the centromere-specific histone H3 (CENH3) (Ahmadli et al., 2023; Jin et al., 2023; Khaitova et al., 2023). We asked if the centromere-mediated mechanism safeguarding genome stability also works under UV. To this end, a weak *cenh3* allele, which shows indistinguishable development and male fertility as wild-type but is sensitive to high temperature (Wang et al., 2023), was used for UV treatment assay. At control condition, around 19.84% metaphase I meiocytes in *cenh3* showed univalents (Fig. 3K and Q), suggesting that the obligate CO formation is lost in these cells; meanwhile, chromosome entanglement was visualized (Fig. 3L). We found that UV-C significantly increased the rate of univalent-harboring meiocytes in *cenh3* (Fig. 3Q), indicating that CO formation is more prone to be attenuated when the centromere is destabilized. Remarkably, *cenh3* exhibited significantly elevated rates of meiocytes at metaphase I to II showing premature segregation of sister chromatids upon UV-C or UV-B radiation, higher than that in Col under the same UV spectrum (Fig. 3K, M and R). Moreover, about 42.50% meiocytes at anaphase I in *cenh3* at control condition displayed chromosome bridges (Fig. 3N, normal; O, abnormal; and S), which indicates an irregularity in segregation of homologous chromosomes. Interestingly, radiation by either UV ray lowered the proportion of meiocytes with chromosome bridges in *cenh3* (Fig. 3S).

To provide further evidence that UV-C perturbs sister chromatid segregation, we performed co-immunolocalization of ɑ-tubulin and CENH3 in Col and *cenh3*. In control, Col showed one and two spindles at metaphase I and II, respectively, in which microtubule filaments got attached to the CENH3-labeled kinetochores (Fig. 3T and U). In UV-C-radiated Col, we observed metaphase II meiocytes with irregularly segregated sister chromatids (Fig. 3V; yellow arrows), indicating a lesion in sister cohesion protection or release. In *cenh3*, the attenuated function of CENH3 did not alter the spindle morphology at metaphase I and/or II (Fig. 3W and X) at both normal and UV-C conditions, but no CENH3 foci was detected likely owing to the deletion of 27 amino acids at the N-terminal tail of CENH3 in this allele (Fig. 3W-A’) (Wang et al., 2023). In UV-C-radiated *cenh3*, metaphase II meiocytes showing asynchronous separation of sister chromatids were observed (Fig. 3Y-A’; yellow arrows). These data suggested that the centromeres may facilitate meiotic chromosome stability in Arabidopsis under UV stress.

### UV reduces CO formation and increases homolog synapsis lesions in *cenh3*

To identify the cause of univalents in UV-radiated *cenh3*, we analyzed crossover formation by examining the chiasma structure, which marks the physical connections between homologous chromosomes during homologous recombination, in Col and *cenh3* at both control and UV conditions. The number of chiasmata can be quantified referring to the morphology of a bivalent (Fig. 4A) (Huang et al., 2022). Col at normal condition generated an average of 9.00 chiasmata per diakinesis meiocyte, which revealed a significant reduction under either UV spectrum (Fig. 4B), suggesting that UV reduces CO formation in Arabidopsis. The negative impact was also detected in *cenh3* (Fig. 4B). However, *cenh3* showed higher scores of chiasmata number than Col at normal and UV-A conditions (Fig. 4B). The similar numbers of chiasmata in Col and *cenh3* under UV-C and UV-B thus hinted higher levels of CO reduction (Fig. 4B).

**Figure 4.**
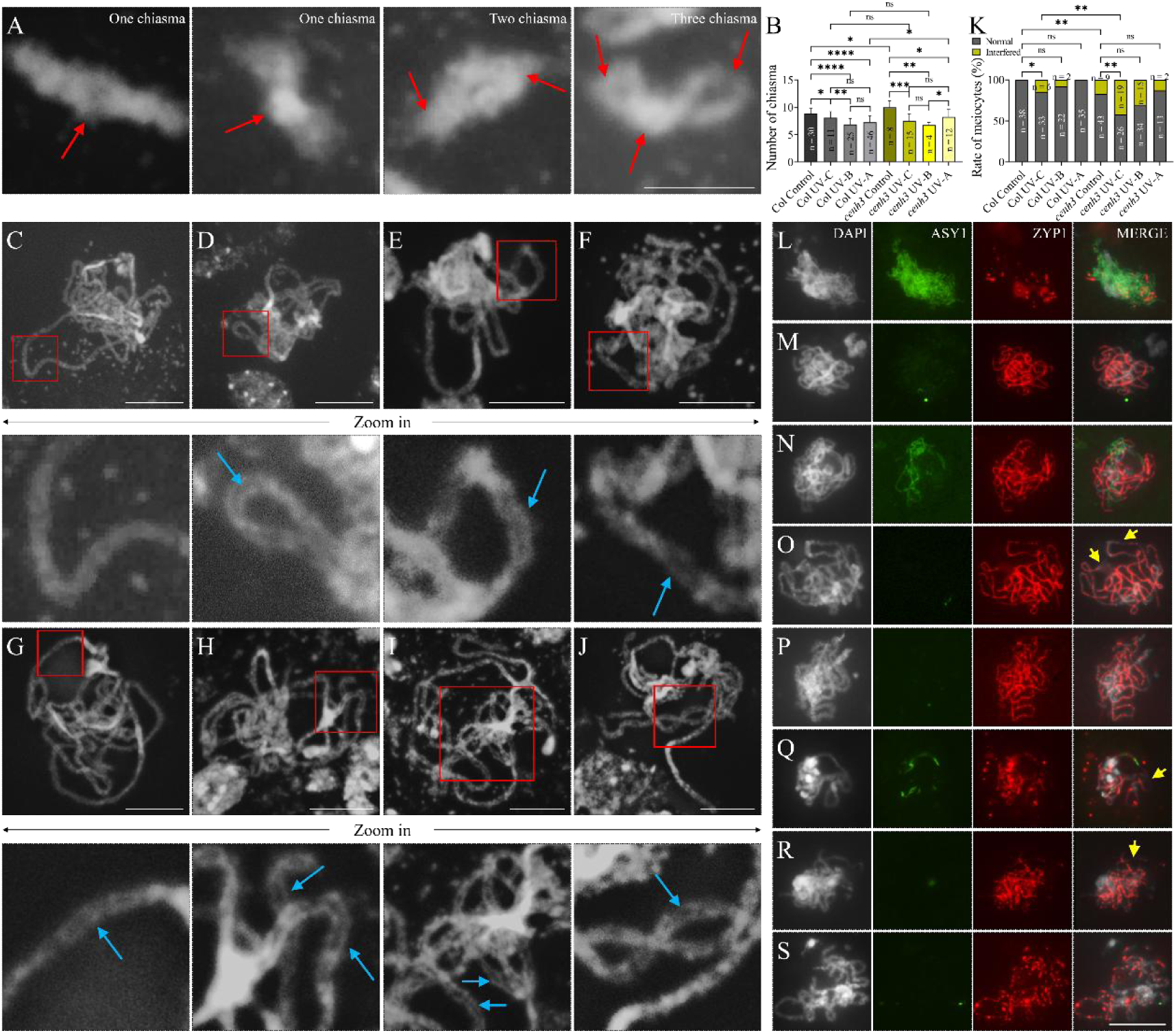
UV reduces CO rate and interferes with homolog synapsis. A, Representative images of DAPI-stained bivalents with one, two or three chiasmata. Red arrows indicate the chiasma location. B, Graph showing the average number of chiasmata per diakinesis meiocyte in Col and *cenh3* at control and different UV conditions; unpaired *t*-tests were performed. C-J, DAPI-stained pachytene chromosome in Col (C and D) and *cenh3* (E-J) at control (C, E and F) and UV-C radiation (G-J) conditions; the zoomed-in figures show the regions labeled by red boxes. K, Graph showing the rate of pachytene meiocytes in Col and *cenh3* at control and different UV conditions showing normal or interfered configurations; χ^2^-tests were performed; n indicates the number of the analyzed cells; **** indicates *P* < 0.0001; *** indicates *P* < 0.001; ** indicates *P* < 0.01; * indicates *P* < 0.05; ns indicates *P* > 0.05. L-S, Co-immunolocalization of ASY1 and ZYP1 on zygotene (L) and pachytene (M-S) chromosomes in Col (L-N) and *cenh3* (O-S) at control (L and M, O and P) and UV-C radiation (N, Q-S) conditions. Yellow arrows indicate chromosome regions with failed assembly of ZYP1. Scale bars in A and the other figures equal 5 and 10 μm, respectively.

Synapsis of homologous chromosomes plays a vital role in maintaining obligate CO formation (France et al., 2021; Yang et al., 2022). Hence, to search for the insight of UV-induced reduction of CO number, we analyzed the morphology of pachytene chromosomes in Col and *cenh3*. In Col at normal condition, homologous chromosomes fully synapsed at pachytene and showed alignment at whole chromosome regions (Fig. 4C). Upon UV-C radiation, the rate of pachytene meiocytes showing co-alignment yet at a greater distance at certain regions between homologous chromosomes was significantly increased, which, however, did not occur under UV-B and UV-A (Fig. 4D and K; blue arrow). The enlarged distance implied that synapsis may be interfered by UV-C. Notably, at control condition, about 17.31% meiocytes in *cenh3* showed the altered chromosome configuration like the effect of UV-C (Fig. 4E, F and K; blue arrows). In addition, the rate significantly increased to 42.22% under UV-C, even higher than that in UV-C-radiated Col (Fig. 4G-K; blue arrows). UV-B and UV-C radiation did not further perturb the proportion of meiocytes with the synapsis lesion (Fig. 4K).

To verify that UV-C interferes with synapsis, we performed co-immunolocalization of ASY1 and ZYP1 to monitor the assembly of synaptonemal complex (SC). In control, ASY1 exhibited a linear configuration and located at the whole regions along the chromosomes axe at zygotene, when ZYP1 started to be loaded at the central region of SC (Fig. 4L). Thereafter at pachytene, ASY1 was removed and ZYP1 was fully assembled, indicating the completion of synapsis (Fig. 4M). In comparison, some pachytene-like meiocytes in UV-C-radiated Col showed a delayed removement of ASY1 signals and incomplete ZYP1 installation on the chromosomes (Fig. 4N), implying that the programmed dynamic behaviors of ASY1 and ZYP1 are altered (Yang et al., 2020; Yang et al., 2022). In *cenh3* at normal condition, incomplete or discontinuous assembly of ZYP1 on pachytene-like chromosomes, on which ASY1 was fully unloaded, was visualized (Fig. 4O and P; yellow arrows), suggesting that *cenh3* has a defect in SC stability. Upon UV-C radiation, more severe ZYP1 configuration occurred in *cenh3*, which exhibited a delayed ASY1 removal, and broken and dotted ZYP1 loading (Fig. 4Q-S; yellow arrows). Therefore, these findings demonstrated that UV-C promotes synapsis instability in *cenh3*.

### UV alters the orientation and organization of spindle and phragmoplast at meiosis II

Chromosome segregation and nuclei positioning require normal assembly and organization of spindles and phragmoplasts. To address how UV radiation interferes with the distribution of the chromosome sets or nuclei at the end of meiosis II, we performed co-immunolocalization of ɑ-tubulin and CENH3 to examine the microtubule organization and the behaviors of centromeres during meiosis. In control, microtubules were organized as a nest-like appearance surrounding chromosomes at prophase I, which progressively became shorter and more concentrated around the chromosomes and were then shaped into a spindle structure at metaphase I (Fig. 5A). At anaphase I, a phragmoplast was formed between the separated homologous chromosomes, which was then spatially guided by the polar microtubule bundles to facilitate the expansion of a temporal cell plate at interkinesis (Fig. 5A). At metaphase II, two spindles were formed to drive the segregation of sister chromatids; and, at telophase II, two phragmoplast structures were built, which were organized into mini-phragmoplasts composed of radial microtubule arrays (RMAs) between the four separated nuclei at tetrad stage (Fig. 5A).

**Figure 5.**
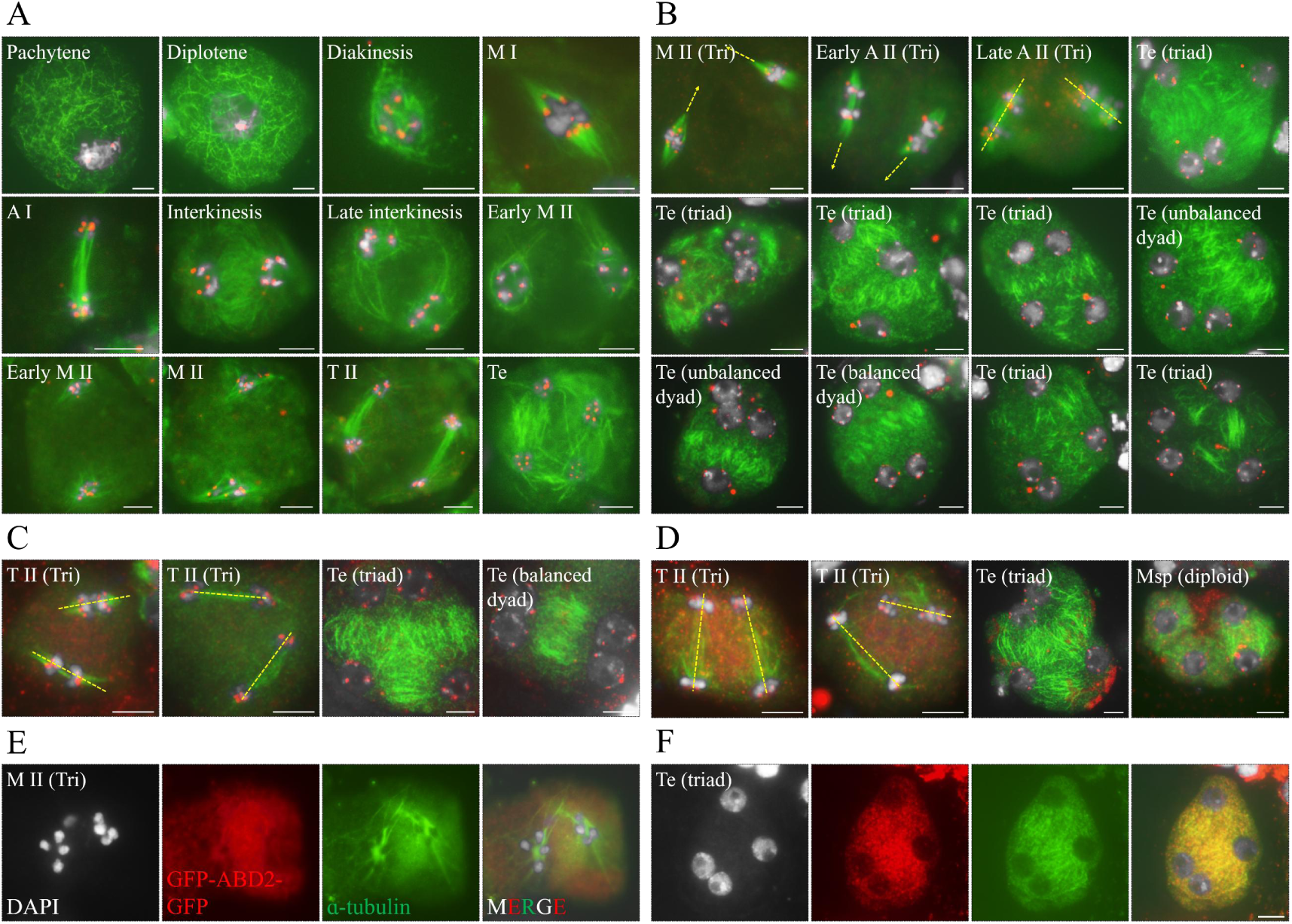
UV radiation interferes with spindle and phragmoplast orientation and radial microtubule array organization during meiosis II. A-D, Co-immunolocalization of ɑ-tubulin and CENH3 in meiocytes of Col at control condition (A), or radiated by UV-C (B), UV-B (C) or UV-A (D). E and F, Co-immunolocalization of GFP-ABD2-GFP and ɑ-tubulin in metaphase II (E) and tetrad (F) meiocytes of Col expressing *pUBQ10::GFP-ABD2-GFP* reporter upon UV-C radiation. Developmental stages are indicated in each figure. M I or II, metaphase I or II; A I or II, anaphase I or II; T II, telophase II; Te, tetrad; Msp, microspore; Tri, triangle configuration. Yellow lines highlight the directions of spindle or phragmoplast. White, DAPI; green, ɑ-tubulin; red, CENH3 or GFP. Scale bars = 5 μm.

In Col radiated by UV-C, UV-B or UV-A, we did not find any obvious difference in microtubule organization during meiosis I (Supplemental Fig. S4). However, spindles at metaphase II displayed altered orientation, leading to arrangement of phragmoplasts with triangle or parallel configurations at anaphase II and/or telophase II (Fig. 5B, C and D, yellow lines; Supplemental Fig. S5; Supplemental Fig. S6A and B). At tetrad stage, haploid nuclei displayed adjacent or clustered distribution, between which microtubule filament showed reduced content, or the assembly of RMAs was omitted, leading to triads and balanced or unbalanced dyads (Fig. 5B, C and D; Supplemental Fig. S5; Supplemental Fig. S6C-J). Moreover, in UV-C-radiated Col, we observed meiocytes with failed assembly of RMA between separated haploid nuclei (Supplemental Fig. S5, blue arrows in the blue star-labeled cells; Supplemental Fig. S6D-F), and meiocytes showing curved RMA filaments (Supplemental Fig. S5; yellow star-labeled cells). These data unveiled that UV interferes with the assembly and organization of microtubule cytoskeleton in meiosis II.

Furthermore, we followed dynamic organization of actin filament in meiosis by monitoring the localization of GFP-tagged fimbrin Actin Binding Domain 2 (ABD2) in Col expressing the *pUBQ10::GFP-ABD2-GFP* reporter using an anti-GFP antibody (Dyachok et al., 2014). The recombinant GFP-ABD2-GFP protein co-localized with microtubules throughout meiosis (Supplemental Fig. S7), which suggested that actin and microtubule cytoskeleton act together to facilitate the dynamic behaviors of chromosomes. After UV-C radiation, the organization of actin cytoskeleton network during meiosis I was not influenced (Supplemental Fig. S8A-D). However, meiocytes with a triangle-like configuration of actin-composed spindles at metaphase II, and triads, balanced and unbalanced dyads at tetrad stage were observed (Fig. 5E and F; Supplemental Fig. S8E-J). In addition, the accumulation of GFP-ABD2-GFP proteins in the meiocytes of UV-C-radiated plants exhibited dotted or a more diffused appearance (Fig. 5F; Supplemental Fig. S8), which suggested an interfered actin stability.

### MAP65-3 localization at tetrad stage is compromised by UV radiation

To further characterize the UV-induced microtubule defects, we monitored the localization of the microtubule-associated protein MAP65-3, which cross-links antiparallel microtubules near their plus ends, using a *pMAP65-3::GFP-MAP65-3* reporter line (Ho et al., 2012; Sofroni et al., 2020). In control, GFP-MAP65-3 localized together with microtubules from prophase I to metaphase I (Supplemental Fig. S9). At interkinesis, GFP-MAP65-3 accumulated at the midzone between the separated nuclei; and, at metaphase II, it co-localized with microtubules to build the spindles (Fig. 6A and B). At anaphase II and tetrad stages, GFP-MAP65-3 were loaded at the midzones between the separated two sets of the sister chromatids and the four haploid nuclei, respectively (Fig. 6C and D).

**Figure 6.**
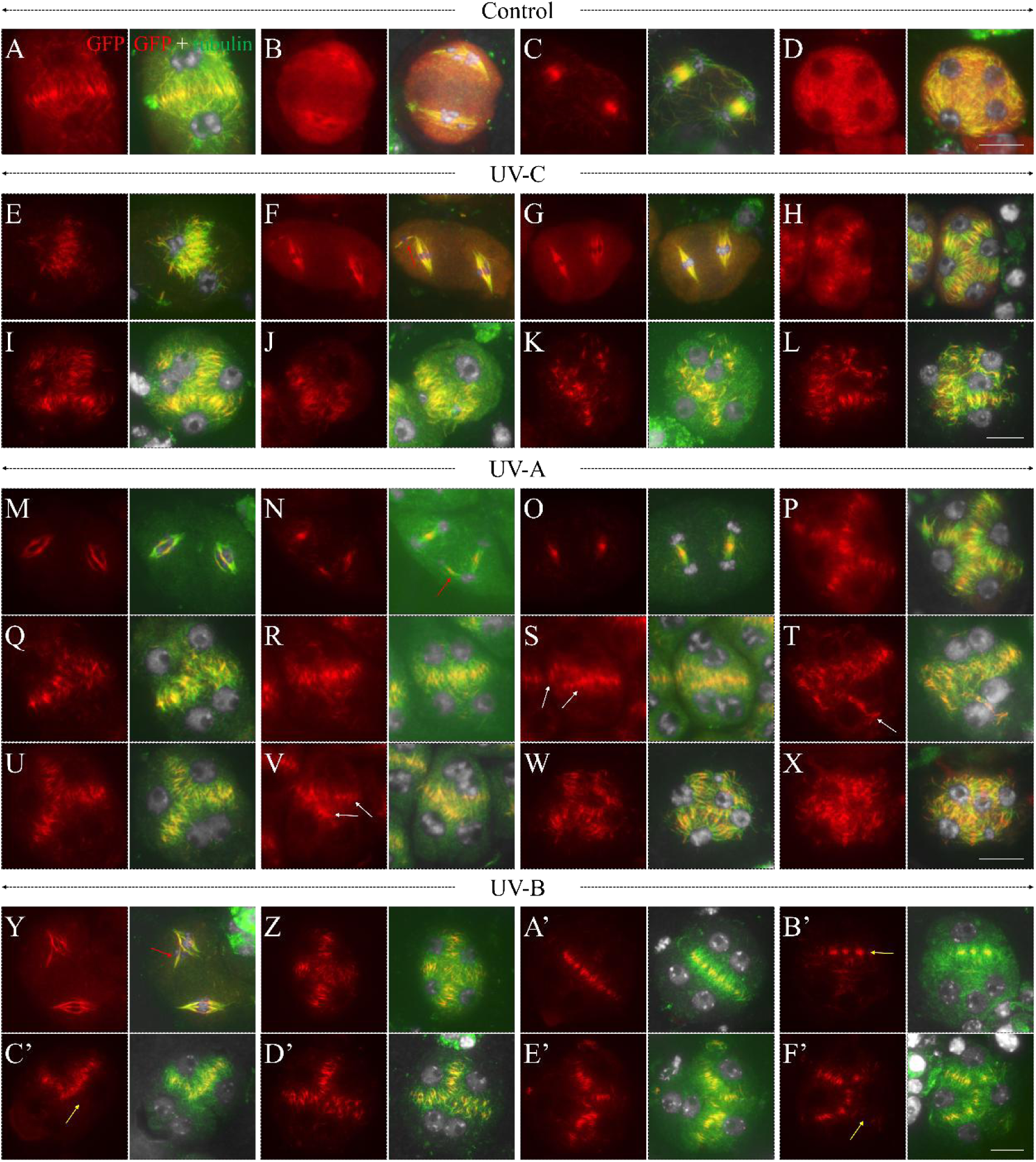
UV interferes with MAP65-3 localization at tetrad stage in Arabidopsis meiosis. A-F’, Co-localization of GFP-MAP65-3 and ɑ-tubulin in interkinesis (A and E), metaphase II (B, F and G, M and Y), telophase II (C, N and O) and tetrad (D, H-L, P-X, and Z-F’) meiocytes in Col at control condition (A-D), or radiated by UV-C (E-L), UV-A (M-X) or UV-B (Y-F’). Red arrows indicate extra assembly of spindles or phragmoplast; white arrows indicate multi-layer or extra accumulation of GFP-MAP65-3; yellow arrows indicate omitted or reduced GFP-MAP65-3 localization. Red, GFP-MAP65-3; green, ɑ-tubulin. Scale bars = 10 μm.

Radiation by either UV spectrum did not alter the microtubule organization and the localization pattern of GFP-MAP65-3 during meiosis I (Supplemental Fig. S10; Supplemental Fig. S11; Supplemental Fig. S12). In UV-C-radiated reporter line, we occasionally observed irregular distribution of phragmoplast between the separated nuclei at interkinesis (Fig. 6E). More obvious defects occurred during meiosis II. At metaphase II and telophase II, both three UV spectrum induced irregular orientation of spindles and/or phragmoplasts (Fig. 6G, M, N, O and Y; Supplemental Fig. S13G, M, N, O and Y), although the localization pattern of GFP-MAP65-3 exhibited the same as in control (Fig. 6B and C, control; F and G, UV-C; M-O, UV-A; Y, UV-B). In addition, extra assembly of spindle or phragmoplast were observed (Fig. 6F, N and Y), which indicated unbalanced chromosome segregation and thus a consequent production of aneuploids (Fig. 6W and X; Supplemental Fig. S10; Supplemental Fig. S13W and X). At tetrad stage, in UV-induced triads, balanced or unbalanced dyads, failed localization of GFP-MAP65-3 between adjacent nuclei (Fig. 6H, P and Z, normal; I-K, Q-V, A’-E’, abnormal; Supplemental Fig. S10; Supplemental Fig. S11; Supplemental Fig. S12), or even the separated nuclei (Fig. 6C’ and F’, yellow arrows), was observed. Meanwhile, some meiocytes showed reduced, sparse, or discontinuous GFP-MAP65-3 content (Fig. 6K, Q, B’ and D’-F’). Moreover, multi-layer or extra accumulation of GFP-MAP65-3 were also visualized (Fig. 6S, T and V, white arrows). Collectively, these data suggested that UV specifically interferes with MAP65-3 localization at tetrad stage.

### UV-induced meiotic restitution is not mediated by the impacted tapetum and UVR8

Interfered or aborted pollen development upon UV radiation suggested that the development or function of tapetum, the most inner cell layer in the anthers, may be perturbed or impaired (Fig. 1A and B; Supplemental Fig. S1). We have previously reported that the Aborted Microspores (AMS)- and Tapetal Development and Function 1 (TDF1)-mediated early development of tapetum is required for normal RMA organization and meiotic cytokinesis (Tidy et al., 2022). Thus, we questioned if UV radiation induces meiotic restitution via the impacted tapetum. To this end, we first examined the expression of AMS in the anthers of Col after UV radiation by performing live-imaging of a *pAMS::AMS-GFP* reporter (Xiong et al., 2016). At normal growth condition, AMS was specifically expressed in the tapetal cell layer, with strong GFP signals being observed at early microsporogenesis stages (Fig. 7A), which decreased at bicellular or tricellular microspore stages due to programmed cell death of the tapetal cells (Supplemental Fig. S14A) (Ferguson et al., 2017; Xiong et al., 2016).

**Figure 7.**
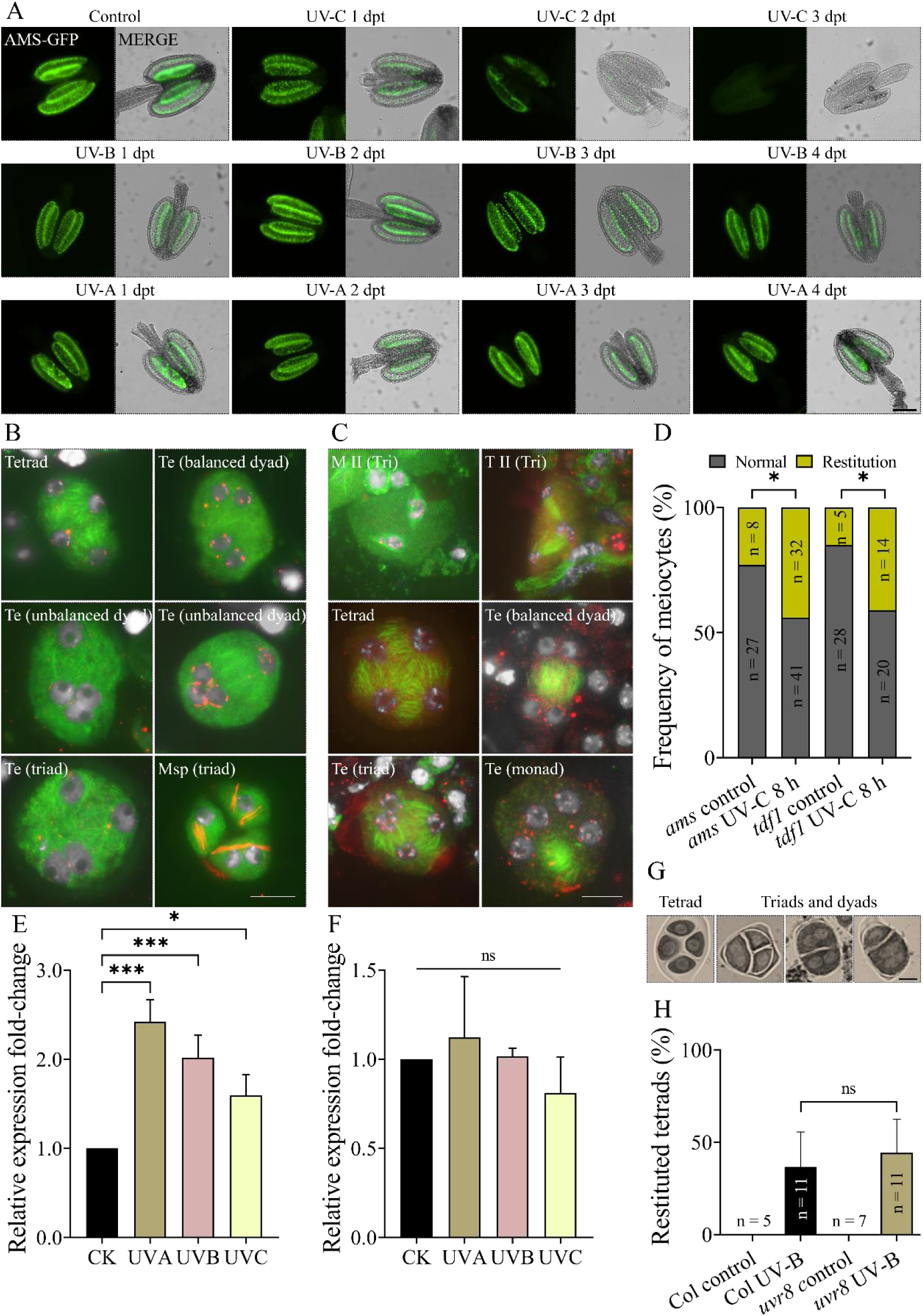
UV increases meiotic restitution rate in *ams* and *tdf1*. A, Live-imaging of *pAMS::AMS-GFP* expression in meiosis-staged anthers of Col at control condition, and at following days post radiation by UV-C for 8 h, and by UV-B or UV-A for 24 h. B, Co-immunolocalization of ɑ-tubulin and CENH3 in meiocytes of Col at 2-4 dpt of UV-C radiation. C, Representative images of co-immunolocalization of ɑ-tubulin and CENH3 in meiocytes of *ams* or *tdf1* at normal condition or at one day post UV-C radiation. D, Graph showing the rates of meiotic restitution in *ams* and *tdf1* at normal condition or at 1 dpt of UV-C radiation. E and F, Relative expression fold-changes of *UVR8* (E) and *COP1* (F) in meiosis-staged flower buds from Col grown at normal condition, or radiated by UV-C for 8 h, or by UV-B or UV-A for 24 h. G, Representative images of orcein-stained meiotic products in *uvr8* at 1 dpt of UV-B radiation. H, Graph showing the rates of meiotically-restituted tetrads in *uvr8* at control and 1 dpt of UV-B radiation. For D, χ^2^-tests were performed; for E and F, one-way ANOVA tests were performed; for H, unpaired *t*-tests were performed; variation bars indicate standard errors; n indicates the number of the analyzed cells or plant individuals; *** indicates *P* < 0.001; * indicates *P* < 0.05. Te, tetrad stage; Msp, microspore; Tri, triangle configuration. White, DAPI; green, ɑ-tubulin; red, CENH3. For A, scale bar = 100 μm; for B, C and G, scale bars = 10 μm.

No alteration in AMS-GFP expression was detected right upon the completion of UV radiation (Supplemental Fig. S15). However, in UV-C-radiated plants, increasingly reduced AMS-GFP expression in both meiosis- and microspore-staged anthers was observed at 1 and 2 dpt; and, at 3 dpt, no AMS-GFP signal can be detected in the anthers (Fig. 7A; Supplemental Fig. S14B). Immunolocalization of ɑ-tubulin in tetrad-staged meiocytes at 2-4 dpt of UV-C treatment showed defective RMA organization and meiotic restitution (Fig. 7B). These data reveal that UV-C impairs tapetum. In comparison, radiation by UV-B or UV-A for 24 h did not affect AMS-GFP expression in the anthers throughout 1-4 dpt (Fig. 7A; Supplemental Fig. S14C and D), which suggested that UV-B and UV-A do not damage tapetum.

The distinct impacts of UV-C and UV-B/A on AMS expression imply that UV interferes with RMA organization and induces meiotic restitution may not by influencing tapetum. To verify our speculation, we analyzed the microtubule organization in meiocytes of *ams* and *dyt1*, which have impaired tapetum function and cytokinesis defects at normal growth condition (Tidy et al., 2022), upon UV-C radiation. To exclude the potential secondary effect by the impact of UV-C on tapetum, only the flower buds at 1 dpt were used in the assay. At control condition, irregular spindle and phragmoplast orientation and meiotically-restituted RMA formation were observed at metaphase II and tetrad stages, respectively, in both *ams* and *tdf1* (Fig. 7C and D). In UV-C-radiated *ams* and *tdf1*, the proportion of meiocytes showing meiotic restitution was significantly increased (Fig. 7D), which supported the hypothesis that UV-induced meiotic restitution is not mediated by the impacted tapetum development.

In Arabidopsis, sensing of UV-B is conducted by UV Resistance Locus 8 (UVR8) (Tilbrook et al., 2013). We applied quantitative reverse transcription PCR (RT-qPCR) to examine the impact of UV on the expression of *UVR8* in the meiosis-staged flower buds. A significantly elevated transcription of *UVR8* was detected upon radiation by either UV spectrum (Fig. 7E), indicating that UVR8 is not only responsive to UV-B. The expression of *Constitutive Photomorphogenic 1* (*COP1*), which interacts with UV-B-activated UVR8 to initiate coordinated response to UV-B (Favory et al., 2009; Huang et al., 2014; Wang et al., 2022b; Yin et al., 2016; Zhang et al., 2023), was not influenced by any UV spectrum (Fig. 7F). To address if UV-induced meiotic restitution is channeled by UVR8, tetrad formation in UV-B-radiated *uvr8* was analyzed, which revealed that *uvr8* underwent normal meiosis at control condition (Fig. 7G and H) and produced a similar level of meiotically-restituted tetrads as Col upon UV-B radiation (Fig. 7G and H). Therefore, UV interferes with meiosis not likely relying on the UVR8-mediated UV perception.

### UV radiation does not further perturb spindle orientation in *jason*

Since UV causes triangle or parallel configurations of spindle and/or phragmoplast at meiosis II (Fig. 5B, C and D) and primarily induces FDR-type unreduced gametes (Fig. 2G and H), we hypothesized that UV-induced meiotic restitution may be owing to an attenuated function of JASON and/or Parallel Spindle 1 (AtPS1), which regulates metaphase II spindle orientation, with both the mutants generating FDR-type diploid gametes (Brownfield et al., 2015; d’Erfurth et al., 2008; De Storme and Geelen, 2011). The expression of *JASON* and *AtPS1* were examined by RT-qPCR, which revealed a non-significantly altered transcription of *JASON*, but a reduction of *AtPS1* upon UV-C and UV-A radiation (Fig. 8A). Moreover, we found that the expression of *MPK4*, which is required for cell plate formation in mitotic and meiotic cytokinesis (Takahashi et al., 2010; Zeng et al., 2011), was upregulated by UV-C and UV-A (Fig. 8A). In comparison, the expression of *OSD1* and *TAM*, which regulate meiotic cell cycle transition (Bulankova et al., 2010; Cromer et al., 2012; d’Erfurth et al., 2010; d’Erfurth et al., 2009), and *ATM*, which plays roles in both DSB repair and meiosis progression under stress (De Jaeger-Braet et al., 2022; Kurzbauer et al., 2021), were not altered by UV radiation (Fig. 8A).

**Figure 8.**
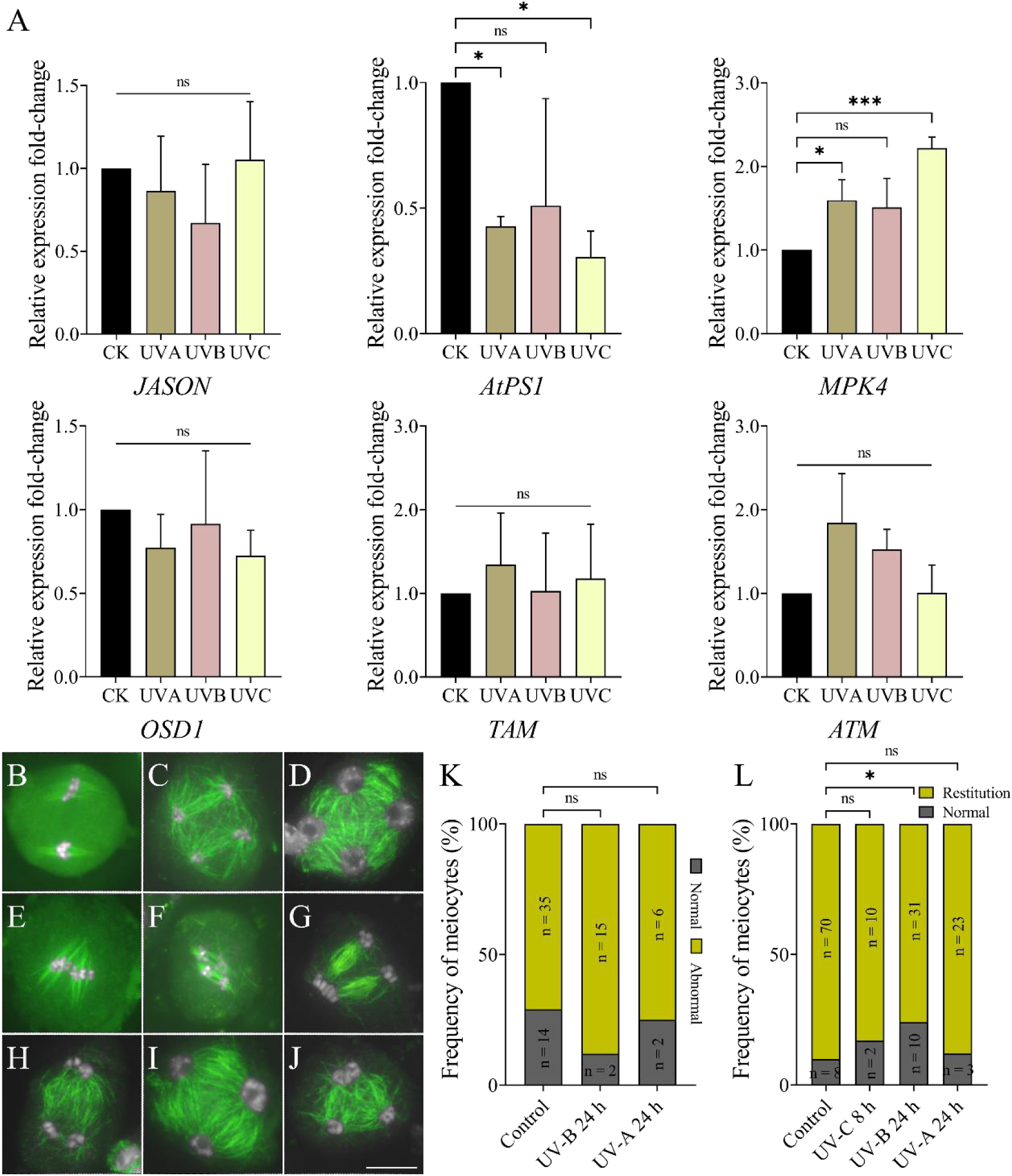
UV induces meiotic restitution in *uvr8*, but does not further perturb spindle orientation in *jason*. A, The relative expression fold-changes of *JASON*, *AtPS1*, *MPK4*, *OSD1*, *TAM* and *ATM* in meiosis-staged flower buds from Col grown at normal condition, or radiated by UV-C for 8 h, or by UV-B or UV-A for 24 h. B-J, Representative images of immunolocalization of ɑ-tubulin in metaphase II (B, E and F), telophase II (C, G and H) and tetrad (D, I and J) meiocytes of *jason* at control condition or radiated by UV, showing normal (B-D) or abnormal (E-J) configurations. K, Graph showing the rate meiocytes showing normal and abnormal spindle or phragmoplast orientation at metaphase II or telophase II. L, Graph showing the rate meiocytes showing normal or meiotically-restituted RMA organization at tetrad stage. For A, one-way ANOVA tests were performed; for K and L, χ^2^-tests were performed; variation bars indicate standard errors; n indicates the number of the analyzed cells; *** indicates *P* < 0.001; * indicates *P* < 0.05; ns indicates *P* > 0.05. Scale bars = 10 μm.

Although UV does not affect the expression of *JASON*, the altered spindle and phragmoplast orientation and the reduced *AtPAS1* expression may still be owing to the compromised JASON function (De Storme and Geelen, 2011). To verify this hypothesis, we analyzed the impact of UV radiation on the microtubule organization at meiosis II in *jason*. At control condition, 71.43% meiocytes in *jason* at metaphase II or telophase II displayed defective orientation of spindle or phragmoplast (Fig. 8B and C, normal spindle and phragmoplast; E-H, triangle and parallel spindle or phragmoplast), which, however, was not significantly changed upon UV-B or UV-A radiation (Fig. 8K). In addition, 89.74% tetrad-staged meiocytes in *jason* at control showed a triad appearance of RMA formation associated with a fused or adjacent positioning of two haploid nuclei (Fig. 8D, normal RMA; I and J, triad-like RMA), which was not altered by UV-C or UV-A but was increased by UV-B (Fig. 8L). These data suggested that UV does not further disturb the spindle and phragmoplast orientation in *jason*, but UV-B might increase meiotic restitution level in *jason* via a different cellular mechanism.

Dysfunction of the PHD domain protein Male Meiocyte Death 1 (MMD1) causes alterations in spindle and phragmoplast orientation at meiosis II, consequently leading to meiotic restitution in Arabidopsis (Andreuzza et al., 2015; Liu et al., 2021). To address if UV induces meiotic restitution partially by affecting MMD1, we generated a *pMMD1::GUS-GFP* reporter and analyzed the impact of UV on the activity of *MMD1* promoter by performing GUS staining. The *MMD1* promoter was specifically active in the meiocytes (Supplemental Fig. S16); and, radiation by either UV spectrum did not obviously alter the color of the anthers (Supplemental Fig. S16), which suggested that UV-induced meiotic restitution is not caused by the alteration in *MMD1* expression.

### UV does not increase meiotic restitution rate in autotetraploid Arabidopsis

In Arabidopsis, meiotic restitution in *jason* can be partially rescued by increased meiocyte size (Wang et al., 2023). We questioned if meiocyte size is also a determinant of UV-induced meiotic restitution. To this end, we analyzed the impact of UV on meiosis in the synthetic autotetraploid Col, in which meiosis is unstable and a low level of meiotic restitution occurs (Fu et al., 2021). At control condition, about 27.27% pachytene meiocytes in autotetraploid Col showed compromised pairing of homologous chromosomes (Fig. 9A, fully paired; B, partially paired; and Q). Unbalanced chromosome segregation was observed at anaphase I, metaphase II and telophase II stages (Fig. 9C, E and H, balanced; D, G and J, unbalanced); meanwhile, chromosomes at metaphase II with a tripolar configuration were visualized, which suggested irregular orientation of spindles (Fig. 9F, yellow arrows). At tetrad stage, both meiotically-restituted products and aneuploids were observed (Fig. 9K, normal; I and L, triad; M, polyad).

**Figure 9.**
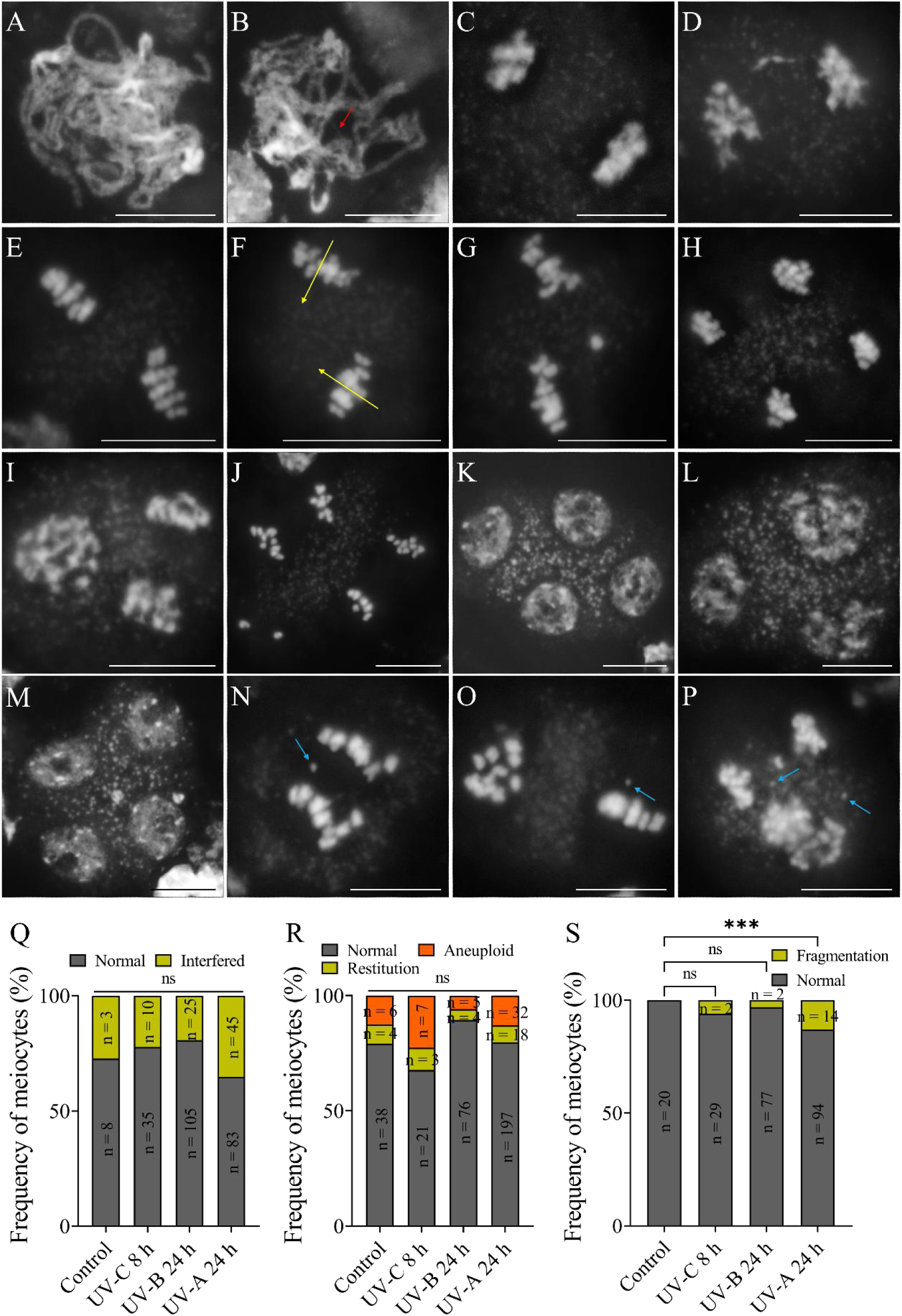
UV does not increase meiotic restitution rate in autotetraploid Col. A-M, Representative images of DAPI-stained pachytene (A and B), anaphase I (C and D), metaphase II (E-G), telophase II (H-J) and tetrad (K-M) meiocytes in autotetraploid Col at control condition or radiated by UV. The red arrow indicates unpaired homologous chromosomes; yellow arrows indicate spindle directions. N-P, DAPI-stained anaphase I (N), metaphase II (O) and telophase II (P) meiocytes showing chromosome fragments in UV-A-radiated autotetraploid Col. Blue arrows indicate chromosome fragments. Q, Graph showing the rate of pachytene meiocytes with normal or interfered chromosome pairing. R, Graph showing the frequency of tetrad meiocytes showing normal configuration, meiotic restitution, or aneuploid nuclei. S, Graph showing the rate of anaphase I, metaphase II and telophase II meiocytes showing normal or interfered chromosome integrity. χ^2^-tests were performed; n indicates the number of the analyzed cells; *** indicates *P* < 0.001; ns indicates *P* > 0.05. Scale bars = 10 μm.

In the autotetraploid Col radiated by either UV-C, UV-B or UV-A, the frequencies of pachytene meiocytes with defects in pairing of homologous chromosomes were not significantly altered (Fig. 9Q), suggesting that UV does not further attenuate synapsis of homologous chromosomes in autotetraploid Arabidopsis. At tetrad stage, both kinds of UV spectrum did not alter the rates of the meiocytes that manifested either meiotic restitution or aneuploids (Fig. 9L, M and R). However, UV-A-radiated autotetraploid Col showed a significantly increased rate of meiocytes harboring small chromosome bodies, which indicate chromosome fragments (Fig. 9N-P, blue arrows; and S) (Kurzbauer et al., 2021; Zhao et al., 2023). Overall, these findings revealed that UV does not induce meiotic restitution in tetraploid Arabidopsis with an enlarged meiocyte size.

### UV induces meiotic restitution in both dicot and monocot plants

To test if UV can induce meiotic restitution in other plant species, we analyzed tetrad formation in diploid *Petunia hybrida* L. (W115) upon UV-C radiation. In control, W115 showed low but detectable rates of restituted meiotic products, including triads, balanced or unbalanced dyads and monads, and polyads (Fig. 10A and C), which indicated that meiosis in petunia is unstable. Upon UV-C radiation, W115 showed a significantly increased rate of meiotic restitution; while, the rate of meiocytes that indicate aneuploid spores, was not altered (Fig. 10B and C). Notably, we observed anaphase I meiocytes with a dyad appearance and abnormal cell wall formation phenotypically mimicking premature meiosis termination (Wijnker et al., 2019), which implied that UV-C may impact meiosis progression in petunia.

**Figure 10.**
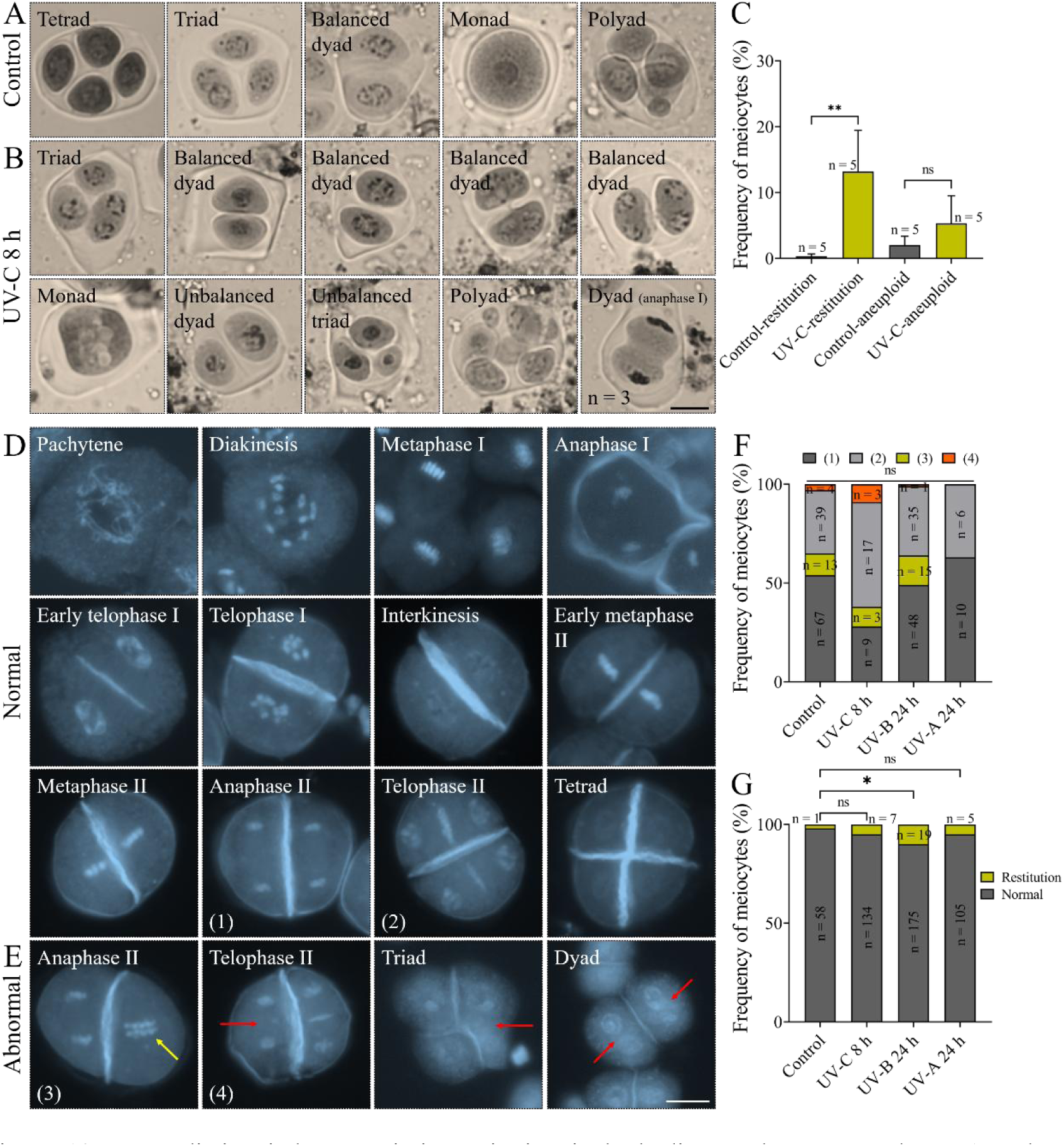
UV radiation induces meiotic restitution in both dicot and monocot plants. A and B, Tetrad-staged meiocytes showing different configurations in diploid *Petunia hybrida* at control condition (A) or radiated by UV-C (B). C, Graph showing the frequencies of meiotic restitution and aneuploid meiotic products in diploid *Petunia hybrida* at control condition or radiated by UV-C; unpaired *t*-tests were performed. D and E, Representative images of combined DAPI and aniline blue staining of meiocytes at different meiosis stages in rice. Yellow arrow indicates unseparated sister chromosomes. Red arrows indicate omission of cell walls. F, Graph showing the frequencies of normal and abnormal chromosome set positioning or cell wall formation in telophase II meiocytes. G, Graph showing the frequencies of normal and meiotically-restituted meiocytes at tetrad stage; χ^2^-tests were performed. n indicates the number of the analyzed cells; ** indicates *P* < 0.01; * indicates *P* < 0.05; ns indicates *P* > 0.05. Scale bars = 10 μm.

Next, we examined the impact of UV radiation on meiosis in rice (*O. sativa* L.) using combined DAPI and aniline blue staining on meiocytes. As a monocot, rice undergoes successive-type of meiotic cytokinesis, in which a cell wall is built following each round of nuclei division (Zhang et al., 2018). In control, a callosic cell wall was initiated at early telophase I from the mid-zone of the cell between the two newly-formed nuclei, and completely extended along the cell plate at interkinesis, leading to production of a dyad (Fig. 10D). After anaphase II, the formation of the second cell wall started at the mid-zone between the separated sister chromatids, and were completely assembled at tetrad stage (Fig. 10D). Occasionally, adjacent chromosome sets, or failed initiation or formation of the second cell wall at anaphase II or telophase II were observed (Fig. 10E, yellow and red arrows). Upon UV-C, UV-B or UV-A radiation, the rates of lesions in chromosome set positioning or cell wall formation at telophase II were not increased (Fig. 10F). However, at tetrad stage, the rate of meiocytes representing restitution was minorly but significantly increased upon UV-B radiation (Fig. 10G). Collectively, these findings suggested that UV-induced meiotic restitution and unreduced gamete formation may broadly occur in flowering plants.

## Discussion

In this study, we showed that UV radiation reduces CO formation and attenuates the stability of centromere-mediated synapsis and sister-chromatid cohesion during meiosis in Arabidopsis. In addition, we showed that UV induces meiotic restitution and unreduced gamete formation in both dicots and monocots. Our findings provide evidence demonstrating that UV has an impact on meiotic genome stability and gametophytic ploidy consistency, which highlights a potential influence of UV on the genome evolution in flowering plants.

In mammals, UV-C-induced DNA lesions CPDs and 6-4PPs can be converted into DSBs due to dysfunctional nucleotide excision repair (NER)-caused aborted DNA replication (Garinis et al., 2005; Oh et al., 2011; Thoma, 1999; Wakasugi et al., 2014). In the present study, however, we showed that UV does not cause defects in meiotic chromosome integrity in both wild-type and mutants defective for DSB repair Arabidopsis (Fig. 3J), which suggest that UV radiation, at least in our experimental conditions, does not induce ectopic DSB formation in the meiocytes. It is possible that UV affects chromosome integrity in a cell-type- or dosage-dependent manner. In UV-A-radiated autotetraploid Arabidopsis, however, the level of chromosome fragmentation is moderately but significantly increased (Fig. 9). We speculate that UV-A may induce specific chromosome-toxic stimulus, which further perturbs chromosomes stability in the autotetraploid Arabidopsis, in which male meiosis is more sensitive to abiotic stresses than the diploids (Fu et al., 2021; Urushibara et al., 2014; Xue et al., 2022).

We detected a reduced chiasmata number in Arabidopsis radiated by either UV spectrum (Fig. 4). It is possible that UV attenuates CO formation by disturbing the activity and/or loading of recombinases, which are sensitive to abiotic stresses (Modliszewski et al., 2018). We found that the obligate bivalent formation is lost in the Arabidopsis *cenh3* mutant, whereas the average number of chiasmata per meiocyte is higher than that in wild-type (Fig. 4). These data suggest that centromere may facilitate CO assurance but, meanwhile, play a role in the restriction of global CO level, likely due to the high density of heterochromatin and the high level of DNA methylation (Naish et al., 2021). We showed that *cenh3* has a defect in synapsis of homologous chromosomes, which may partially contribute to the loss of CO assurance and the higher CO level via the release of chromosome tension and attenuated CO interference (Capilla-Pérez et al., 2021; France et al., 2021; Yang et al., 2022).

UV-C induces a low rate of meiocytes with premature separation of sister chromatids in wild-type, and this effect is pronounced in *cenh3* (Fig. 3), in addition, the synapsis lesions in *cenh3* under UV is more severe (Fig. 4). Thus, the centromeres play a role in stabilizing SC structure and sister-chromatid cohesion under UV stress. We speculate that UV may further attenuate the centromere-mediated cohesion complex stability (García-Nieto et al., 2023). It is also possible that the irregular sister-chromatids separation takes place in the UV-induced univalents, in which sister kinetochores undergo irregular orientation in meiosis I (Chan and Cande, 1998; Hunt et al., 1995; Khodjakov et al., 1997; Rebollo and Arana, 1995; Xue et al., 2019; Yamamoto and Hiraoka, 2003). In yeast, the rates of recombination between either intersister-chromatids or homologous chromosomes are elevated by UV-induced DNA damages, and the release of sister-chromatid cohesion promotes this effect (Covo et al., 2010). However, this mechanism might not be conserved between species since we found that the CO rates in both wild-type and *cenh3* is reduced upon UV radiation (Fig. 4). Considering that functional centromeres are required for maintenance of chromosome integrity under heat stress (Ahmadli et al., 2023; Jin et al., 2023; Wang et al., 2023), it may be a common feature that the centromeres protect meiotic genome stability in plants under diverse stresses. Nevertheless, under UV condition, *cenh3* shows a similar chiasmata number as that in Col (Fig. 4), the centromeres thus probably do not mediate the stability of meiotic recombination under UV stress.

On the other hand, *cenh3* at control condition showed chromosome bridges at anaphase I (Fig. 3), likely due to the reduced efficiency or the lesion in the microtubule-kinetochore interaction (Capitao et al., 2021). In insects, UV radiation on kinetochores at anaphase I in spermatocytes delays or inhibits the movement of homologous chromosomes to cell polars (Ilagan and Forer, 1997; Wong and Forer, 2004). Similar effect has been recorded in UV-C-radiated mitotic cells in mammals (Zhang et al., 2013). Specifically, Zhang et al. reported that UV-C enhances the recruitment of the spindle checkpoint proteins MPS1 and MAD2 and the Aurora B kinase, which activates the spindle assembly checkpoint (SAC) and thus delays the segregation of homologous chromosomes (Zhang et al., 2013). We speculate that the effect of UV-C on SAC activation might be conserved between mitosis and meiosis and across species, which explains the significantly rescued abnormal chromosome interactions in UV-C-radiated *cenh3* (Fig. 3).

UV alters nuclei positioning at the end of meiosis II, which leads to production of unreduced gametes primarily derived from the first meiotic cell division restitution (Fig. 2). Similarly, in UV-radiated rice, no obvious lesion was observed in meiosis I, and defective meiotic cell wall formation only occurred after the second nuclei division (Fig. 10). These findings suggest that the processes involved in ensuring ploidy-halving during meiosis II in plants are especially sensitive to UV radiation. In Arabidopsis, both cold and heat stresses induce meiotic restitution by predominantly targeting the RMA formation after telophase II (De Storme et al., 2012; De Storme and Geelen, 2020; Liu et al., 2018). Hence, the stage during meiosis II when spindles and phragmoplast are organized may be a special time-window for the environmental factors to impose impacts on the ploidy stability of gametes in flowering plants.

Either UV spectrum does not affect the morphology of microtubule cytoskeleton in meiosis I, but specifically alters the orientation of spindle and phragmoplast, which result in omitted or compromised organization of RMA at late meiosis II with resultant meiotically-restituted dyads and triads (Fig. 5; Supplemental Fig. S5; Supplemental Fig. S6; Supplemental Fig. S7; Supplemental Fig. S8). We speculate that UV-C, UV-B and UV-A induce meiotic restitution at least partially via the same cellular mechanism: by interfering with the orientation and organization of spindles and phragmoplasts in meiosis II, which are crucial for spacing the separated chromosome sets and cytokinesis (Brownfield et al., 2015). We found that UV does not further increase the rate of meiocytes with altered spindle and phragmoplast orientation in *jason* (Fig. 8), supporting a hypothesis that UV induces meiotic restitution at least partially by affecting JASON function. However, RT-qPCR analysis showed that UV does not affect the expression of *JASON*, but lowers the expression of *AtPS1* (Fig. 8). We speculate that UV may attenuate JASON function not at the transcription level but by disturbing its subcellular localization; and, the downregulated *AtPS1* expression could be owing to the compromised JASON function (Fig. 8) (De Storme and Geelen, 2011).

The microtubule-associated protein MAP65-3 does not localize between the fused and/or adjacent haploid nuclei at tetrad stage in the UV-induced triads and dyads (Fig. 6), suggesting that the UV-induced RMA organization defects may be at least partially caused by the lesions in MAP65-3-dependent cell plate formation (Ho et al., 2012). However, UV does not affect the localization pattern of MAP65-3 throughout meiosis prior to tetrad stage, even at metaphase II and telophase II stages when the orientation of spindles and phragmoplasts are altered (Fig. 6; Supplemental Fig. S10; Supplemental Fig. S11; Supplemental Fig. S12; Supplemental Fig. S13). The lesions in MAP65-3 localization thus are not likely resulted from the perturbed function the meiotic cell cycle regulator Cyclin Dependent Kinase A;1 (CDKA;1), which regulates microtubule cytoskeleton and cytokinesis with the mutant exhibiting premature MAP65-3-mediated cell plate formation in prophase I (Sofroni et al., 2020). The completely omitted loading, or reduced signals, or discontinuous accumulation (Fig. 6) suggest that the expression or stability of MAP65-3, which are sensitive to abiotic stresses (Liang et al., 2023; Ma and Liu, 2019; Zhang et al., 2012), may be interfered by UV. Moreover, the defective accumulation of MAP65-3 may not be the secondary effect of the adjacent nuclei positioning, since we observed meiocytes with completely abolished MAP65-3 localization between the isolated nuclei, which have certain space in between (Fig. 6).

On the other hand, UV may interfere with spindle and phragmoplast organization and RMA formation through a direct impact on the synthesis and/or stability of microtubules (Grzanka et al., 2006; Jiang et al., 2011; Misovic et al., 2013). In support of this, we observed the occurrence of meiotic restitution upon short periods of UV-C radiation (Supplemental Fig. S2), which indicates a fast response of meiosis to UV stimulus. Meanwhile, the localization of GFP-ABD2-GFP in UV-C-radiated meiocytes exhibits dotted and/or more diffused appearances (Fig. 5; Supplemental Fig. S8), implying an interfered stability of actin. Moreover, UV can affect plant development by inducing ROS, which, in mammals, disturbs the polymerization and depolymerization of actin and its binding proteins (de Jager et al., 2017; Ishimoto and Mori, 2022; Xu et al., 2019; Xue et al., 2022; Yokawa and Baluška, 2015). The irregularities of GFP-ABD2-GFP localization in UV-radiated plants thus may be partially caused by the ROS-caused instability of actin.

In addition to the primary impact on cytoskeleton organization in meiosis II, UV may induce meiotic restitution by affecting one or more processes during meiosis I. This notion could be favored by the following observations: 1), we detected one thirds of unreduced gametes as the SDR-type (Fig. 2); and 2), the slightly but significantly increased rate of meiotically-restituted products in *jason* upon UV-B radiation (Fig. 8) implies that partial UV-induced triads and dyads result from cellular alterations in JASON-independent mechanisms. Because we did not find meiocytes with an obviously arrested or aborted meiosis II in the UV-radiated Col, UV-induced SDR-type unreduced gametes in Arabidopsis are not likely caused by the dysfunction of CYCA1;2/TAM and/or Omission of Second Division (OSD1), two factors controlling meiosis progression and cell cycle transition (d’Erfurth et al., 2010), whose expression is UV-insensitive (Fig. 8). However, in UV-radiated petunia, the dyads, which phenotypically mimic a premature termination of meiosis at telophase I (Fig. 10), suggest that different mechanisms may underpin the UV-induced meiotic restitution between species.

How are UV signals transduced into the meiocytes? We hypothesized that UV-induced meiotic alterations are caused by the impacted tapetum, because: 1), both UV spectrum interfere with gametogenesis especially pollen wall formation (Fig. 1; Supplemental Fig. S1), which largely relies on normal tapetum function (Alonso-Peral et al., 2010; Ferguson et al., 2017; Lou et al., 2018); 2), the early development of tapetum is required for programmed meiosis progression and successful RMA organization and cytokinesis in Arabidopsis (Lei and Liu, 2020; Tidy et al., 2022). However, this hypothesis was undermined by the observations that UV-C impairs AMS expression in the tapetum, but UV-B and UV-A do not (Fig. 7), which reveals distinct impacts of different UV spectrum on the tapetum; and, UV-C increases meiotic restitution rate in *ams* and *tdf1* (Fig. 7). These findings suggest that UV interferes with cytoskeleton organization and induces meiotic restitution independently of the tapetum function, but possibly via a direct impact on the meiocytes. Furthermore, *uvr8* produces a similar frequency of triads and dyads as Col upon UV-B (Fig. 7), we thus conclude that UV-induced meiotic restitution does not rely on the UVR8-mediated UV perception. It is of a particular interest to decipher how UV is sensed and transduced into meiocytes and interferes with the meiotic stability in plants.

## Materials and methods

### Plant materials and growth conditions

Arabidopsis (*Arabidopsis thaliana*) ecotype Columbia (Col-0) was used as the wild-type. The *qrt* (Francis et al., 2006), *atm-2* (Kurzbauer et al., 2021), *atr* (Wang et al., 2021a), *rfc1* (Yu et al., 2012), *xip-101* (Fan et al., 2022), *ku70* (Riha et al., 2002), *sog1* (Wang et al., 2022a), *tdf1* (Tidy et al., 2022), *ams* (Tidy et al., 2022), *jason* (Yi et al., 2023), *cenh3-8* (Wang et al., 2023) and *uvr8* (Huang et al., 2014) mutants, the fluorescent-tagged line (FTL) I3bc (Francis et al., 2007), *pCENH3::RFP-CENH3* (Komaki et al., 2020), *pUBQ10::GFP-ABD2-GFP* (Dyachok et al., 2014), *pMAP65-3::GFP-MAP65-3* (Sofroni et al., 2020) and the *pAMS::AMS-GFP* (Xiong et al., 2016) reporter lines were used in this study. The *pMMD1::GUS-GFP* reporter line was newly generated. Seeds were germinated in 1/2 MS medium for 8 d, and seedlings were transferred to soil and cultivated in a growth chamber with a 16 h day/8 h night and 20℃ condition. The rice (*O. sativa* L.) cultivar ‘Kitaake’ were grown during the growing season in Wuhan (30.52°N, 114.31°E); the *Petunia hybrida* L. (Petunia × hybrida) cultivar ‘Mitchell Diploid’ (W115) were grown in a growth chamber with a 16 h day/8 h night and 20℃ condition.

### Treatment of ultraviolet radiation

Young flowering plants were transferred to a growth chamber with a 20°C and dark condition, and were irradiated with UV-C (λ = 254.30 nm, irradiance *E* = 4.37 W/m^2^), UV-B (λ = 313.00 nm, irradiance *E* = 2.25 W/m^2^), or UV-A (λ = 365.70 nm, irradiance *E* = 3.40 W/m^2^) for different durations.

### Cytological analysis of meiocytes and microspores

Alexander staining of anthers, which were collected from plants in control or at different time points after UV radiation, was performed as described by (Alexander, 1969). Orcein staining, and the combined staining by DAPI and aniline were performed by releasing meiotic products in a drop of 4.5% lactopropionic orcein solution, or in a drop of mixture composed of DAPI (5 μg/mL) and aniline blue solution (0.1% in 0.033% K_3_PO_4_), respectively.

### Pollen germination assay

Flower buds carrying mature pollen grains from control plants or at 6 or 7 dpt of UV radiation was tipped on a glass slide with pollen growth medium (5 mM CaCl_2_, 1 mM MgSO_4_, 5 mM KCl, 0.01 mM H_3_BO_3_, 18% sucrose, 1.5% agarose, pH at 7.5 modified by 0.1 M NaOH). After 8 h incubation under dark in a humid chamber, pollen tubes were observed under a microscope.

### β-glucuronidase (GUS) staining

To analyze the impact of UV radiation or impaired tapetum on the expression of *MMD1*, flower buds harboring meiosis-staged meiocytes from Col plants expressing the *pMMD1::GUS-GFP* reporter were isolated for histochemical GUS staining using a GUS staining kit (Real-Times, RTU4032) following the manufacturer’s protocol.

### Preparation of chromosome spreads

Inflorescences of young, flowering Arabidopsis plants were collected and fixed in pre-cooled Carnoy’s fixative for at least 24 h. Meiosis-staged flower buds were washed twice with distilled water and once with citrate buffer (10 mM, pH = 4.5), followed by incubation in a digestion enzyme mixture (0.3% pectolyase and 0.3% cellulase in citrate buffer, 10 mM, pH = 4.5) at 37°C for 2.5 h. Subsequently, the digested flower buds were washed once in distilled water and macerated in 3 μL distilled water on a glass slide. Two aliquots of 10 μL 60% acetic acid were added to the slide, which was dried on a hotplate at 45°C for 2 min. The slide was washed with 200 μL ice-cold Carnoy’s fixative and then air-dried. Next, 8 μL DAPI (5 μg/mL) diluted in Vectashield antifade mounting medium (Vector Laboratories, https://vectorlabs.com/) was added to the slide, and the coverslip was mounted and sealed with nail polish.

### Protein immunolocalization

Immunolocalization assays were performed by referring to (Liu et al., 2017; Wang et al., 2014). Antibodies against CENH3 (rabbit) (Fu et al., 2021) and ɑ-tubulin (rat) (Lei et al., 2020) were diluted 1:500; antibody against GFP (rabbit) (Zhao et al., 2023) were diluted 1:300; antibodies against SYN1 (mouse) (Zhao et al., 2023), ASY1 (mouse) (Zhao et al., 2023), ZYP1 (rabbit) (Ning et al., 2021), DMC1 (rabbit) (Ning et al., 2021) and ɤH2A.X (rabbit) (Zhao et al., 2023) were diluted 1:100. The secondary antibodies; i.e., goat anti-rabbit Alexa Fluor 555, goat anti-rabbit Alexa Fluor Plus 488, goat anti-mouse Alexa Fluor Plus 488 and F(ab’)2-goat anti-mouse Alexa Fluor Plus 555 from Thermo Fisher Scientific, were diluted to 5 µg/mL.

### Generation of the *pMMD1::GUS-GFP* reporter

To generate the *pMMD1::GUS-GFP* reporter, promoter sequences were selected using RegSite Plant DB, Softberry database and TSSPlant, a tool developed to predict RNA polymerase II transcription start sites in plant DNA sequences (Shahmuradov et al., 2017). The promoter of *MMD1* was amplified using specific primer proAt1g66170. The promoter was cloned into the entry vector pDONR221 and fused to eGFP-GUS in vector pKGWFS7.0. The used primers are listed in Supplemental Table S1. The transformation was performed via the floral dip method (Clough and Bent, 1998) using *Agrobacterium tumefaciens* (GV3101).

### Quantitative reverse transcription PCR

Total RNA was isolated from meiosis-staged flower buds of Arabidopsis Col plants grown at normal conditions and radiated by UV-C for 8 h, or by UV-B or UV-A for 24 h, respectively. The first-strand cDNA was synthesized using a FastKing cDNA kit (TIANGEN), and RT-qPCR was performed on a quantitative PCR instrument (Bio-Rad) using SYBR qPCR Master Mix (Vazyme) according to manufacturer’s instructions. For each gene at both the control and UV radiation conditions, three biological replicates and three technical replicates were performed. *Actin7* (AT5G09810) was used as the reference gene, and the relative expression levels of target genes were calculated using the comparative CT method. Significance was calculated using the one-way ANOVA test and significance level was set as *P* < 0.05. The used primers are listed in Supplemental Table S1.

### Live-imaging of reporters

The expression of RFP-CENH3 or RFP in FTL line I3c was monitored by releasing microspores or pollen grains in a drop of distilled water, and were then visualized under a fluorescence microscope. To analyze the expression of AMS in the anthers, anthers at different stages in flowering Col plants expressing the *pAMS::AMS-GFP* reporter were isolated and placed on a glass slide with a drop of distilled water being added to the samples, which was then mounted with a cover slide and examined under an inverted fluorescence microscope. The developmental stage was determined by checking the meiocytes or microspores in the anthers.

### Microscopy

Images were captured by an Olympus IX83 inverted fluorescence microscope with an X-Cite lamp and a Prime BSI camera, or an M-Shot ML31 microscope equipped with an MS60 camera. Bifluorescent images and Z-stacks were processed using ImageJ. The brightness and contrast setting of pictures were adjusted using PowerPoint 2016.

### Statistical analysis

Significance analysis was performed using unpaired *t* tests or Chi-squared tests with GraphPad Prism (version 8) software. The numbers of analyzed cells or the biological replicates are shown in the figures or legends; variation bars indicate standard deviations.

## Acknowledgements

The authors thank Arp Schnittger (Universität Hamburg) for sharing the *pMAP65-3::MAP65-3-GFP* and *pCENH3::TagRFP-CENH3* reporters. They appreciate Elison B. Blancaflor (NRI) for sharing the *pUBQ10::GFP-ABD2-GFP* reporter. They thank Yingxiang Wang (SCAU) for sharing the *rfc1* mutant. They thank Shunping Yan (HZAU) for providing the *atr*, *atm* and the *ku70* mutants. They thank Hongtao Liu (CAS) for sharing the *uvr8* mutant. They thank Jiang Hua (IPK) for sharing the *jason* mutant. They thank Zhongnan Yang, Jun Zhu and Yue Lou (SHNU) for sharing the *tdf1* and *ams* mutants and the *pAMS::AMS-GFP* reporter. They thank Gregory Copenhaver (UNC) for providing the FTL line I3bc. They thank Jianzhong Lin (Hunan University) for providing the rice seeds.

## Author contributions

H.F., J.Z., J.Z., L.H., C.W., D.M., W.P., L.S. and Z.R. performed investigation; T.F., Z.W., W.W., X.L., G.Y., J.L., Y.Z. and D.G. contributed to data analysis; B.L. conceived project, analyzed data, wrote, and edited manuscript. All the authors have read the manuscript and approved the submission.

## Supplemental data

Supplemental Figure S1. UV-C radiation disrupts pollen development in *Arabidopsis thaliana*.

Supplemental Figure S2. Short periods of UV-C radiation induce meiotic restitution in *Arabidopsis thaliana*.

Supplemental Figure S3. UV-C does not obviously affect ɤH2A.X and DMC1 foci formation on prophase I chromosomes.

Supplemental Figure S4. UV does not affect assembly and organization of microtubule during meiosis I.

Supplemental Figure S5. UV induces irregular organization of spindle and phragmoplast during meiosis II in Col

Supplemental Figure S6. UV-C induces irregular organization of spindle and phragmoplast during meiosis II in Col.

Supplemental Figure S7. Organization of actin cytoskeleton in meiocytes of *Arabidopsis thaliana*.

Supplemental Figure S8. UV-C radiation interferes with the organization of actin- and tubulin-mediated phragmoplast during meiosis II in *Arabidopsis thaliana*.

Supplemental Figure S9. Co-immunolocalization of GFP-MAP65-3 and ɑ-tubulin in meiosis I meiocytes in *Arabidopsis thaliana*.

Supplemental Figure S10. UV-C affects the organization of MAP65-3 and microtubules at tetrad stage but not during meiosis I in Col.

Supplemental Figure S11. UV-A affects the organization of MAP65-3 and microtubules at tetrad stage but not during meiosis I in Col.

Supplemental Figure S12. UV-B affects the organization of MAP65-3 and microtubules at tetrad stage but not during meiosis I in Col.

Supplemental Figure S13. UV interferes with MAP65-3 localization at tetrad stage in *Arabidopsis thaliana*.

Supplemental Figure S14. UV-C, but not UV-B or UV-A, impairs AMS expression in the anthers.

Supplemental Figure S15. AMS expression in the anthers of Col is not affected right upon the finish of UV radiation.

Supplemental Figure S16. UV does not influence the activity of *MMD1* promoter. Supplemental Table S1. Primers used in this study.

## Data availability

All data and materials that support the findings of this study are available upon request from the corresponding author.

## References

1. Ahmadli, U., Kalidass, M., Khaitova, L.C., Fuchs, J., Cuacos, M., Demidov, D., Zuo, S., Pecinkova, J., Mascher, M., Ingouff, M., Heckmann, S., Houben, A., Riha, K., and Lermontova, I. (2023). High temperature increases centromere-mediated genome elimination frequency and enhances haploid induction in Arabidopsis. Plant Communications 4, 100507.

2. Alexander, M.P. (1969). Differential staining of aborted and nonaborted pollen. Stain Technol 44, 117–122.

3. Alonso-Peral, M.M., Li, J., Li, Y., Allen, R.S., Schnippenkoetter, W., Ohms, S., White, R.G., and Millar, A.A. (2010). The microRNA159-regulated GAMYB-like genes inhibit growth and promote programmed cell death in Arabidopsis. Plant Physiology 154, 757–771.

4. Andreuzza, S., Nishal, B., Singh, A., and Siddiqi, I. (2015). The chromatin protein DUET/MMD1 controls expression of the meiotic gene TDM1 during male meiosis in Arabidopsis. Plos Genetics 11, e1005396.

5. Armstrong, S.J., Caryl, A.P., Jones, G.H., and Franklin, F.C. (2002). Asy1, a protein required for meiotic chromosome synapsis, localizes to axis-associated chromatin in Arabidopsis and Brassica. Journal of Cell Science 115, 3645–3655.

6. Barnes, P.W., Williamson, C.E., Lucas, R.M., Robinson, S.A., Madronich, S., Paul, N.D., Bornman, J.F., Bais, A.F., Sulzberger, B., Wilson, S.R., Andrady, A.L., McKenzie, R.L., Neale, P.J., Austin, A.T., Bernhard, G.H., Solomon, K.R., Neale, R.E., Young, P.J., Norval, M., Rhodes, L.E., Hylander, S., Rose, K.C., Longstreth, J., Aucamp, P.J., Ballaré, C.L., Cory, R.M., Flint, S.D., de Gruijl, F.R., Häder, D.-P., Heikkilä, A.M., Jansen, M.A.K., Pandey, K.K., Robson, T.M., Sinclair, C.A., Wängberg, S.-Å., Worrest, R.C., Yazar, S., Young, A.R., and Zepp, R.G. (2019). Ozone depletion, ultraviolet radiation, climate change and prospects for a sustainable future. Nature Sustainability 2, 569–579.

7. Bell, B.A., Fletcher, W.J., Ryan, P., Seddon, A.W.R., Wogelius, R.A., and Ilmen, R. (2018). UV-B-absorbing compounds in modern Cedrus atlantica pollen: the potential for a summer UV-B proxy for Northwest Africa. The Holocene 28, 1382–1394.

8. Benca, J.P., Duijnstee, I.A.P., and Looy, C.V. (2018). UV-B–induced forest sterility: implications of ozone shield failure in Earth’s largest extinction. Science Advances 4, e1700618.

9. Brar, G.A., and Amon, A. (2008). Emerging roles for centromeres in meiosis I chromosome segregation. Nature Reviews Genetics 9, 899–910.

10. Brownfield, L., Yi, J., Jiang, H., Minina, E.A., Twell, D., and Köhler, C. (2015). Organelles maintain spindle position in plant meiosis. Nature Communications 6, 6492.

11. Bulankova, P., Riehs-Kearnan, N., Nowack, M.K., Schnittger, A., and Riha, K. (2010). Meiotic progression in Arabidopsis is governed by complex regulatory interactions between SMG7, TDM1, and the meiosis I–specific cyclin TAM Plant Cell 22, 3791–3803.

12. Burma, S., Chen, B.P., Murphy, M., Kurimasa, A., and Chen, D.J. (2001). ATM phosphorylates histone H2AX in response to DNA double-strand breaks. Journal of Biological Chemistry 276, 42462–42467.

13. Capilla-Pérez, L., Durand, S., Hurel, A., Lian, Q., Chambon, A., Taochy, C., Solier, V., Grelon, M., and Mercier, R. (2021). The synaptonemal complex imposes crossover interference and heterochiasmy in Arabidopsis. Proc Natl Acad Sci U S A 118, e2023613118.

14. Capitao, C., Tanasa, S., Fulnecek, J., Raxwal, V.K., Akimcheva, S., Bulankova, P., Mikulkova, P., Crhak Khaitova, L., Kalidass, M., Lermontova, I., Mittelsten Scheid, O., and Riha, K. (2021). A CENH3 mutation promotes meiotic exit and restores fertility in SMG7-deficient Arabidopsis. Plos Genetics 17, e1009779.

15. Çetinbaş-Genç, A., Toksöz, O., Piccini, C., Kilin, Ö., Sesal, N.C., and Cai, G. (2022). Effects of UV-B radiation on the performance, antioxidant response and protective compounds of Hazelnut pollen. Plants 11, 2574.

16. Chan, A., and Cande, W.Z. (1998). Maize meiotic spindles assemble around chromatin and do not require paired chromosomes. Journal of Cell Science 111, 3507–3515.

17. Clough, S.J., and Bent, A.F. (1998). Floral dip: a simplified method for Agrobacterium-mediated transformation of Arabidopsis thaliana. Plant J 16, 735–743.

18. Covo, S., Westmoreland, J.W., Gordenin, D.A., and Resnick, M.A. (2010). Cohesin is limiting for the suppression of DNA damage–induced recombination between homologous chromosomes. Plos Genetics 6, e1001006.

19. Cromer, L., Heyman, J., Touati, S., Harashima, H., Araou, E., Girard, C., Horlow, C., Wassmann, K., Schnittger, A., De Veylder, L., and Mercier, R. (2012). OSD1 promotes meiotic progression via APC/C inhibition and forms a regulatory network with TDM and CYCA1;2/TAM. Plos Genetics 8, e1002865.

20. d’Erfurth, I., Jolivet, S., Froger, N., Catrice, O., Novatchkova, M., and Mercier, R. (2009). Turning meiosis into mitosis. PLoS Biol 7, e1000124.

21. d’Erfurth, I., Jolivet, S., Froger, N., Catrice, O., Novatchkova, M., Simon, M., Jenczewski, E., and Mercier, R. (2008). Mutations in AtPS1(Arabidopsis thaliana Parallel Spindle 1) lead to the production of diploid pollen grains. Plos Genetics 4.

22. d’Erfurth, I., Cromer, L., Jolivet, S., Girard, C., Horlow, C., Sun, Y., To, J.P.C., Berchowitz, L.E., Copenhaver, G.P., and Mercier, R. (2010). The cyclin-A CYCA1;2/TAM is required for the meiosis I to meiosis II transition and cooperates with OSD1 for the prophase to first meiotic division transition. Plos Genetics 6, e1000989.

23. Da Ines, O., Degroote, F., Goubely, C., Amiard, S., Gallego, M.E., and White, C.I. (2013). Meiotic recombination in Arabidopsis is catalysed by DMC1, with RAD51 playing a supporting role. Plos Genetics 9, e1003787.

24. De Jaeger-Braet, J., Krause, L., Buchholz, A., and Schnittger, A. (2022). Heat stress reveals a specialized variant of the pachytene checkpoint in meiosis of Arabidopsis thaliana. Plant Cell 34, 433–454.

25. de Jager, T.L., Cockrell, A.E., and Du Plessis, S.S. (2017). Ultraviolet light induced generation of reactive oxygen species. Adv Exp Med Biol 996, 15–23.

26. De, S., Jose, J., Pal, A., Roy Choudhury, S., and Roy, S. (2022). Exposure to low UV-B dose induces DNA double-strand breaks mediated onset of dndoreduplication in Vigna radiata (L.) R. Wilczek seedlings. Plant and Cell Physiology 63, 463–483.

27. De Storme, N., and Geelen, D. (2011). The Arabidopsis mutant jason produces unreduced first division restitution male gametes through a parallel/fused spindle mechanism in meiosis II. Plant Physiology 155, 1403–1415.

28. De Storme, N., and Geelen, D. (2020). High temperatures alter cross-over distribution and induce male meiotic restitution in Arabidopsis thaliana. Communications Biology 3, 187.

29. De Storme, N., Copenhaver, G.P., and Geelen, D. (2012). Production of diploid male gametes in Arabidopsis by cold-induced destabilization of postmeiotic radial microtubule arrays. Plant Physiology 160, 1808–1826.

30. De Storme, N., Zamariola, L., Mau, M., Sharbel, T.F., and Geelen, D. (2013). Volume-based pollen size analysis: an advanced method to assess somatic and gametophytic ploidy in flowering plants. Plant Reproduction 26, 65–81.

31. Del Valle, J.C., Buide, M.L., Whittall, J.B., Valladares, F., and Narbona, E. (2020). UV radiation increases phenolic compound protection but decreases reproduction in Silene littorea. Plos One 15, e0231611.

32. Diffey, B.L. (2002). Sources and measurement of ultraviolet radiation. Methods 28, 4–13.

33. Dyachok, J., Sparks, J.A., Liao, F., Wang, Y.-S., and Blancaflor, E.B. (2014). Fluorescent protein-based reporters of the actin cytoskeleton in living plant cells: fluorophore variant, actin binding domain, and promoter considerations. Cytoskeleton 71, 311–327.

34. Fan, T., Kang, H., Wu, D., Zhu, X., Huang, L., Wu, J., and Zhu, Y. (2022). Arabidopsis γ-H2A.X-INTERACTING PROTEIN participates in DNA damage response and safeguards chromatin stability. Nature Communications 13, 7942.

35. Favory, J.J., Stec, A., Gruber, H., Rizzini, L., Oravecz, A., Funk, M., Albert, A., Cloix, C., Jenkins, G.I., Oakeley, E.J., Seidlitz, H.K., Nagy, F., and Ulm, R. (2009). Interaction of COP1 and UVR8 regulates UV-B-induced photomorphogenesis and stress acclimation in Arabidopsis. EMBO Journal 28, 591–601.

36. Feng, H., An, L., Tan, L., Hou, Z., and Wang, X. (2000). Effect of enhanced ultraviolet-B radiation on pollen germination and tube growth of 19 taxa in vitro. Environmental and Experimental Botany 43, 45–53.

37. Ferguson, A.C., Pearce, S., Band, L.R., Yang, C., Ferjentsikova, I., King, J., Yuan, Z., Zhang, D., and Wilson, Z.A. (2017). Biphasic regulation of the transcription factor ABORTED MICROSPORES (AMS) is essential for tapetum and pollen development in Arabidopsis. New Phytologist 213, 778–790.

38. France, M.G., Enderle, J., Röhrig, S., Puchta, H., Franklin, F.C.H., and Higgins, J.D. (2021). ZYP1 is required for obligate cross-over formation and cross-over interference in Arabidopsis. Proc Natl Acad Sci U S A 118.

39. Francis, K.E., Lam, S.Y., and Copenhaver, G.P. (2006). Separation of Arabidopsis pollen tetrads is regulated by QUARTET1, a pectin methylesterase gene. Plant Physiology 142, 1004–1013.

40. Francis, K.E., Lam, S.Y., Harrison, B.D., Bey, A.L., Berchowitz, L.E., and Copenhaver, G.P. (2007). Pollen tetrad-based visual assay for meiotic recombination in Arabidopsis. Proc Natl Acad Sci U S A 104, 3913–3918.

41. Fraser, W.T., Lomax, B.H., Jardine, P.E., Gosling, W.D., and Sephton, M.A. (2014). Pollen and spores as a passive monitor of ultraviolet radiation. Frontiers in Ecology and Evolution 2.

42. Fraser, W.T., Sephton, M.A., Watson, J.S., Self, S., Lomax, B.H., James, D.I., Wellman, C.H., Callaghan, T.V., and Beerling, D.J. (2011). UV-B absorbing pigments in spores: biochemical responses to shade in a high-latitude birch forest and implications for sporopollenin-based proxies of past environmental change. Polar Research 30, 8312.

43. Fu, H., Zhao, J., Ren, Z., Yang, K., Wang, C., Zhang, X., Elesawi, I.E., Zhang, X., Xia, J., Chen, C., Lu, P., Chen, Y., Liu, H., Yu, G., and Liu, B. (2021). Interfered chromosome pairing at high temperature promotes meiotic instability in autotetraploid Arabidopsis. Plant Physiology 188, 1210–1228.

44. García-Nieto, A., Patel, A., Li, Y., Oldenkamp, R., Feletto, L., Graham, J.J., Willems, L., Muir, K.W., Panne, D., and Rowland, B.D. (2023). Structural basis of centromeric cohesion protection. Nat Struct Mol Biol 30, 853–859.

45. Garinis, G.A., Mitchell, J.R., Moorhouse, M.J., Hanada, K., de Waard, H., Vandeputte, D., Jans, J., Brand, K., Smid, M., van der Spek, P.J., Hoeijmakers, J.H., Kanaar, R., and van der Horst, G.T. (2005). Transcriptome analysis reveals cyclobutane pyrimidine dimers as a major source of UV-induced DNA breaks. EMBO Journal 24, 3952–3962.

46. Gill, S.S., Anjum, N.A., Gill, R., Jha, M., and Tuteja, N. (2015). DNA damage and repair in plants under ultraviolet and ionizing radiations. The Scientific World Journal 2015, 250158.

47. Gordon, S.G., and Rog, O. (2023). Building the synaptonemal complex: molecular interactions between the axis and the central region. Plos Genetics 19, e1010822.

48. Grzanka, D., Domaniewski, J., Grzanka, A., and Zuryn, A. (2006). Ultraviolet radiation (UV) induces reorganization of actin cytoskeleton in CHOAA8 cells. Neoplasma 53, 328–332.

49. Hartung, F., Wurz-Wildersinn, R., Fuchs, J., Schubert, I., Suer, S., and Puchta, H. (2007). The catalytically active tyrosine residues of both SPO11-1 and SPO11-2 are required for meiotic double-strand break induction in Arabidopsis. Plant Cell 19, 3090–3099.

50. Hessling, M., Haag, R., Sieber, N., and Vatter, P. (2021). The impact of far-UVC radiation (200-230 nm) on pathogens, cells, skin, and eyes - a collection and analysis of a hundred years of data. GMS Hyg Infect Control 16, Doc07.

51. Higgins, J.D., Sanchez-Moran, E., Armstrong, S.J., Jones, G.H., and Franklin, F.C. (2005). The Arabidopsis synaptonemal complex protein ZYP1 is required for chromosome synapsis and normal fidelity of crossing over. Genes & Development 19, 2488–2500.

52. Ho, C.M., Lee, Y.R., Kiyama, L.D., Dinesh-Kumar, S.P., and Liu, B. (2012). Arabidopsis microtubule-associated protein MAP65-3 cross-links antiparallel microtubules toward their plus ends in the phragmoplast via its distinct C-terminal microtubule binding domain. Plant Cell 24, 2071–2085.

53. Huang, J., Li, X., Wang, C., and Wang, Y. (2022). Evaluation of crossover number, distribution, and interference using cytological assays in Arabidopsis. Curr Protoc 2, e599.

54. Huang, X., Yang, P., Ouyang, X., Chen, L., and Deng, X.W. (2014). Photoactivated UVR8-COP1 module determines photomorphogenic UV-B signaling output in Arabidopsis. Plos Genetics 10, e1004218.

55. Hunt, P., LeMaire, R., Embury, P., Sheean, L., and Mroz, K. (1995). Analysis of chromosome behavior in intact mammalian oocytes: monitoring the segregation of a univalent chromosome during female meiosis. Hum Mol Genet 4, 2007–2012.

56. Ilagan, A.B., and Forer, A. (1997). Effects of ultraviolet-microbeam irradiation of kinetochores in crane-fly spermatocytes. Cell Motil Cytoskeleton 36, 266–275.

57. Ishimoto, T., and Mori, H. (2022). Control of actin polymerization via reactive oxygen species generation using light or radiation. Front Cell Dev Biol 10, 1014008.

58. Jiang, L., Wang, Y., Björn, L.O., and Li, S. (2011). UV-B-induced DNA damage mediates expression changes of cell cycle regulatory genes in Arabidopsis root tips. Planta 233, 831–841.

59. Jin, C., Sun, L., Trinh, H.K., and Danny, G. (2023). Heat stress promotes haploid formation during CENH3-mediated genome elimination in Arabidopsis. Plant Reproduction 36, 147–155.

60. Khaitova, L.C., Mikulkova, P., Pecinkova, J., Kalidass, M., Heckmann, S., Lermontova, I., and Riha, K. (2023). Heat stress impairs centromere structure and segregation of meiotic chromosomes in Arabidopsis. BioRxiv, 2023.2004.2019.537519.

61. Khodjakov, A., Cole, R.W., McEwen, B.F., Buttle, K.F., and Rieder, C.L. (1997). Chromosome fragments possessing only one kinetochore can congress to the spindle equator. Journal of Cell Biology 136, 229–240.

62. Komaki, S., Takeuchi, H., Hamamura, Y., Heese, M., Hashimoto, T., and Schnittger, A. (2020). Functional analysis of the plant chromosomal passenger complex. Plant Physiology 183, 1586–1599.

63. Koti, S., Reddy, K.R., Kakani, V.G., Zhao, D., and Reddy, V.R. (2004). Soybean (Glycine max) pollen germination characteristics, flower and pollen morphology in response to enhanced ultraviolet-B radiation. Ann Bot 94, 855–864.

64. Kurzbauer, M.-T., Janisiw, M.P., Paulin, L.F., Prusén Mota, I., Tomanov, K., Krsicka, O., Haeseler, A.v., Schubert, V., and Schlögelhofer, P. (2021). ATM controls meiotic DNA double-strand break formation and recombination and affects synaptonemal complex organization in plants. Plant Cell.

65. Kurzbauer, M.T., Uanschou, C., Chen, D., and Schlogelhofer, P. (2012). The recombinases DMC1 and RAD51 are functionally and spatially separated during meiosis in Arabidopsis. Plant Cell 24, 2058–2070.

66. Lei, X., and Liu, B. (2020). Tapetum-dependent male meiosis progression in plants: increasing evidence emerges. Frontiers in Plant Science 10, 1667.

67. Lei, X., Ning, Y., Eid Elesawi, I., Yang, K., Chen, C., Wang, C., and Liu, B. (2020). Heat stress interferes with chromosome segregation and cytokinesis during male meiosis in Arabidopsis thaliana. Plant Signaling & Behavior 15, 1746985.

68. Li, J., Yang, L., Jin, D., Nezames, C.D., Terzaghi, W., and Deng, X.W. (2013). UV-B-induced photomorphogenesis in Arabidopsis. Protein Cell 4, 485–492.

69. Li, J., Liang, W., Liu, Y., Ren, Z., Ci, D., Chang, J., and Qian, W. (2022). The Arabidopsis ATR-SOG1 signaling module regulates pleiotropic developmental adjustments in response to 3’-blocked DNA repair intermediates. Plant Cell 34, 852–866.

70. Liang, M., Ji, T., Wang, X., Wang, X., Li, S., Gao, L., Ma, S., and Tian, Y. (2023). Comprehensive analyses of microtubule-associated protein MAP65 family genes in Cucurbitaceae and CsaMAP65s expression profiles in cucumber. Journal of Applied Genetics 64, 393–408.

71. Liu, B., De Storme, N., and Geelen, D. (2017). Cold interferes with male meiotic cytokinesis in Arabidopsis thaliana independently of the AHK2/3-AHP2/3/5 cytokinin signaling module. Cell Biology International 41, 879–889.

72. Liu, B., De Storme, N., and Geelen, D. (2018). Cold-induced male meiotic restitution in Arabidopsis thaliana is not mediated by GA-DELLA signaling. Frontiers in Plant Science 9, 91.

73. Liu, B., Jin, C., De Storme, N., Schotte, S., Schindfessel, C., De Meyer, T., and Geelen, D. (2021). A hypomorphic mutant of PHD domain protein Male Meiocytes Death 1. Genes (Basel) 12, 516.

74. Liu, Q., Wang, J., Miki, D., Xia, R., Yu, W., He, J., Zheng, Z., Zhu, J.-K., and Gong, Z. (2010). DNA Replication Factor C1 mediates genomic stability and transcriptional gene silencing in Arabidopsis Plant Cell 22, 2336–2352.

75. Liu, Y., Deng, Y., Li, G., and Zhao, J. (2013). Replication factor C1 (RFC1) is required for double-strand break repair during meiotic homologous recombination in Arabidopsis. Plant Journal 73, 154–165.

76. Lou, Y., Zhou, H.-S., Han, Y., Zeng, Q.-Y., Zhu, J., and Yang, Z.-N. (2018). Positive regulation of AMS by TDF1 and the formation of a TDF1–AMS complex are required for anther development in Arabidopsis thaliana. New Phytologist 217, 378–391.

77. Ma, H., and Liu, M. (2019). The microtubule cytoskeleton acts as a sensor for stress response signaling in plants. Molecular Biology Reports 46, 5603–5608.

78. Misovic, M., Milenkovic, D., Martinovic, T., Ciric, D., Bumbasirevic, V., and Kravic-Stevovic, T. (2013). Short-term exposure to UV-A, UV-B, and UV-C irradiation induces alteration in cytoskeleton and autophagy in human keratinocytes. Ultrastruct Pathol 37, 241–248.

79. Modliszewski, J.L., Wang, H., Albright, A.R., Lewis, S.M., Bennett, A.R., Huang, J., Ma, H., Wang, Y., and Copenhaver, G.P. (2018). Elevated temperature increases meiotic crossover frequency via the interfering (Type I) pathway in Arabidopsis thaliana. Plos Genetics 14, e1007384.

80. Mullenders, L.H.F. (2018). Solar UV damage to cellular DNA: from mechanisms to biological effects. Photochemical & Photobiological Sciences 17, 1842–1852.

81. Naish, M., Alonge, M., Wlodzimierz, P., Tock, A.J., Abramson, B.W., Schmücker, A., Mandáková, T., Jamge, B., Lambing, C., Kuo, P., Yelina, N., Hartwick, N., Colt, K., Smith, L.M., Ton, J., Kakutani, T., Martienssen, R.A., Schneeberger, K., Lysak, M.A., Berger, F., Bousios, A., Michael, T.P., Schatz, M.C., and Henderson, I.R. (2021). The genetic and epigenetic landscape of the Arabidopsis centromeres. Science 374, eabi7489.

82. Ning, Y., Liu, Q., Wang, C., Qin, E., Wu, Z., Wang, M., Yang, K., Elesawi, I.E., Chen, C., Liu, H., Qin, R., and Liu, B. (2021). Heat stress interferes with formation of double-strand breaks and homolog synapsis. Plant Physiology 185, 1783–1797.

83. Niu, B., Wang, L., Zhang, L., Ren, D., Ren, R., Copenhaver, G.P., Ma, H., and Wang, Y. (2015). Arabidopsis Cell Division Cycle 20.1 is required for normal meiotic spindle assembly and chromosome segregation. Plant Cell 27, 3367–3382.

84. Oh, K.S., Bustin, M., Mazur, S.J., Appella, E., and Kraemer, K.H. (2011). UV-induced histone H2AX phosphorylation and DNA damage related proteins accumulate and persist in nucleotide excision repair-deficient XP-B cells. DNA Repair 10, 5–15.

85. Paull, T.T. (2015). Mechanisms of ATM activation. Annu Rev Biochem 84, 711–738.

86. Peterson, R., Slovin, J.P., and Chen, C. (2010). A simplified method for differential staining of aborted and non-aborted pollen grains. International Journal of Plant Biology 1, e13.

87. Ramsey, J., and Schemske, D.W. (1998). Pathways, mechanisms, and rates of polyploid formation in flowering plants. Annual Review of Ecology and Systematics 29, 467–501.

88. Rastogi, R.P., Richa Kumar, A., Tyagi, M.B., and Sinha, R.P. (2010). Molecular mechanisms of ultraviolet radiation-induced DNA damage and repair. J Nucleic Acids 2010, 592980.

89. Ravi, M., and Chan, S.W.L. (2010). Haploid plants produced by centromere-mediated genome elimination. Nature 464, 615–618.

90. Ravi, M., Marimuthu, M.P.A., and Siddiqi, I. (2008). Gamete formation without meiosis in Arabidopsis. Nature 451, 1121–U1110.

91. Rebollo, E., and Arana, P. (1995). A comparative study of orientation at behavior of univalent in living grasshopper spermatocytes. Chromosoma 104, 56–67.

92. Ren, R., Wang, H.F., Guo, C.C., Zhang, N., Zeng, L.P., Chen, Y.M., Ma, H., and Qi, J. (2018). Widespread whole genome duplications contribute to genome complexity and species diversity in angiosperms. Molecular Plant 11, 414–428.

93. Ries, G., Heller, W., Puchta, H., Sandermann, H., Seidlitz, H.K., and Hohn, B. (2000). Elevated UV-B radiation reduces genome stability in plants. Nature 406, 98–101.

94. Riha, K., Watson, J.M., Parkey, J., and Shippen, D.E. (2002). Telomere length deregulation and enhanced sensitivity to genotoxic stress in Arabidopsis mutants deficient in Ku70. Embo j 21, 2819–2826.

95. Roeles, J., and Tsiavaliaris, G. (2019). Actin-microtubule interplay coordinates spindle assembly in human oocytes. Nature Communications 10, 4651.

96. Sampson, B.J., and Cane, J.H. (1999). Impact of enhanced ultraviolet-B radiation on flower, pollen, and nectar production. American Journal of Botany 86, 108–114.

97. Sanchez-Moran, E., Santos, J.L., Jones, G.H., and Franklin, F.C. (2007). ASY1 mediates AtDMC1-dependent interhomolog recombination during meiosis in Arabidopsis. Genes & Development 21, 2220–2233.

98. Schalk, C., Cognat, V., Graindorge, S., Vincent, T., Voinnet, O., and Molinier, J. (2017). Small RNA-mediated repair of UV-induced DNA lesions by the DNA DAMAGE-BINDING PROTEIN 2 and ARGONAUTE 1. Proc Natl Acad Sci U S A 114, E2965–E2974.

99. Shahmuradov, I.A., Umarov, R.K., and Solovyev, V.V. (2017). TSSPlant: a new tool for prediction of plant Pol II promoters. Nucleic acids research 45, e65–e65.

100. Shi, C., and Liu, H. (2021). How plants protect themselves from ultraviolet-B radiation stress. Plant Physiology 187, 1096–1103.

101. Shiomi, Y., Hayashi, A., Ishii, T., Shinmyozu, K., Nakayama, J., Sugasawa, K., and Nishitani, H. (2012). Two different replication factor C proteins, Ctf18 and RFC1, separately control PCNA-CRL4Cdt2-mediated Cdt1 proteolysis during S phase and following UV irradiation. Molecular Cell Biology 32, 2279–2288.

102. Sinha, R.P., and Häder, D.P. (2002). UV-induced DNA damage and repair: a review. Photochem Photobiol Sci 1, 225–236.

103. Sofroni, K., Takatsuka, H., Yang, C., Dissmeyer, N., Komaki, S., Hamamura, Y., Böttger, L., Umeda, M., and Schnittger, A. (2020). CDKD-dependent activation of CDKA;1 controls microtubule dynamics and cytokinesis during meiosis. Journal of Cell Biology 219.

104. Soltis, P.S., Marchant, D.B., Van de Peer, Y., and Soltis, D.E. (2015). Polyploidy and genome evolution in plants. Current Opinion in Genetics & Development 35, 119–125.

105. Su, H., Cheng, Z., Huang, J., Lin, J., Copenhaver, G.P., Ma, H., and Wang, Y. (2017). Arabidopsis RAD51, RAD51C and XRCC3 proteins form a complex and facilitate RAD51 localization on chromosomes for meiotic recombination. Plos Genetics 13, e1006827.

106. Takahashi, M., Teranishi, M., Ishida, H., Kawasaki, J., Takeuchi, A., Yamaya, T., Watanabe, M., Makino, A., and Hidema, J. (2011). Cyclobutane pyrimidine dimer (CPD) photolyase repairs ultraviolet-B-induced CPDs in rice chloroplast and mitochondrial DNA. Plant Journal 66, 433–442.

107. Takahashi, Y., Soyano, T., Kosetsu, K., Sasabe, M., and Machida, Y. (2010). HINKEL kinesin, ANP MAPKKKs and MKK6/ANQ MAPKK, which phosphorylates and activates MPK4 MAPK, constitute a pathway that is required for cytokinesis in Arabidopsis thaliana. Plant and Cell Physiology 51, 1766–1776.

108. Tamura, K., Adachi, Y., Chiba, K., Oguchi, K., and Takahashi, H. (2002). Identification of Ku70 and Ku80 homologues in Arabidopsis thaliana: evidence for a role in the repair of DNA double-strand breaks. Plant Journal 29, 771–781.

109. Thoma, F. (1999). Light and dark in chromatin repair: repair of UV-induced DNA lesions by photolyase and nucleotide excision repair. EMBO Journal 18, 6585–6598.

110. Tidy, A.C., Ferjentsikova, I., Vizcay-Barrena, G., Liu, B., Yin, W., Higgins, J.D., Xu, J., Zhang, D., Geelen, D., and Wilson, Z.A. (2022). Sporophytic control of pollen meiotic progression is mediated by tapetum expression of ABORTED MICROSPORES. Journal of Experimental Botany 73, 5543–5558.

111. Tilbrook, K., Arongaus, A.B., Binkert, M., Heijde, M., Yin, R., and Ulm, R. (2013). The UVR8 UV-B photoreceptor: perception, signaling and response. Arabidopsis Book 11, e0164.

112. Torabinejad, J., Caldwell, M.M., Flint, S.D., and Durham, S. (1998). Susceptibility of pollen to UV-B radiation: an assay of 34 taxa. American Journal of Botany 85, 360–369.

113. Urushibara, A., Kodama, S., and Yokoya, A. (2014). Induction of genetic instability by transfer of a UV-A-irradiated chromosome. Mutation Research/Genetic Toxicology and Environmental Mutagenesis 766, 29–34.

114. Van de Peer, Y., Ashman, T.-L., Soltis, P.S., and Soltis, D.E. (2020). Polyploidy: an evolutionary and ecological force in stressful times. Plant Cell 33, 11–26.

115. Verdaasdonk, J.S., and Bloom, K. (2011). Centromeres: unique chromatin structures that drive chromosome segregation. Nature Reviews Molecular Cell Biology 12, 320–332.

116. Verdaguer, D., Jansen, M.A.K., Llorens, L., Morales, L.O., and Neugart, S. (2017). UV-A radiation effects on higher plants: exploring the known unknown. Plant Science 255, 72–81.

117. Wakasugi, M., Sasaki, T., Matsumoto, M., Nagaoka, M., Inoue, K., Inobe, M., Horibata, K., Tanaka, K., and Matsunaga, T. (2014). Nucleotide excision repair-dependent DNA double-strand break formation and ATM signaling activation in mammalian quiescent cells. Journal of Biological Chemistry 289, 28730–28737.

118. Wang, L., Zhan, L., Zhao, Y., Huang, Y., Wu, C., Pan, T., Qin, Q., Xu, Y., Deng, Z., Li, J., Hu, H., Xue, S., and Yan, S. (2021a). The ATR-WEE1 kinase module inhibits the MAC complex to regulate replication stress response. Nucleic acids research 49, 1411–1425.

119. Wang, N., Gent, J.I., and Dawe, R.K. (2021b). Haploid induction by a maize cenh3 null mutant. Science Advances 7.

120. Wang, X., Wang, L., Huang, Y., Deng, Z., Li, C., Zhang, J., Zheng, M., and Yan, S. (2022a). A plant-specific module for homologous recombination repair. Proc Natl Acad Sci U S A 119, e2202970119.

121. Wang, Y., and Copenhaver, G.P. (2018). Meiotic recombination: mixing it up in plants. Annual Review of Plant Biology 69, 577–609.

122. Wang, Y., Cheng, Z., Lu, P., Timofejeva, L., and Ma, H. (2014). Molecular cell biology of male meiotic chromosomes and isolation of male meiocytes in Arabidopsis thaliana. Methods Mol Biol 1110, 217–230.

123. Wang, Y., Wang, L., Guan, Z., Chang, H., Ma, L., Shen, C., Qiu, L., Yan, J., Zhang, D., Li, J., Deng, X.W., and Yin, P. (2022b). Structural insight into UV-B–activated UVR8 bound to COP1. Science Advances 8, eabn3337.

124. Wang, Z., Chen, M., Yang, H., Hu, Z., Yu, Y., Xu, H., Yan, S., Yi, K., and Li, J. (2023). A simple and highly efficient strategy to induce both paternal and maternal haploids through temperature manipulation. Nature Plants 9, 699–705.

125. Wei, P., Demulder, M., David, P., Eekhout, T., Yoshiyama, K.O., Nguyen, L., Vercauteren, I., Eeckhout, D., Galle, M., De Jaeger, G., Larsen, P., Audenaert, D., Desnos, T., Nussaume, L., Loris, R., and De Veylder, L. (2021). Arabidopsis casein kinase 2 triggers stem cell exhaustion under Al toxicity and phosphate deficiency through activating the DNA damage response pathway. Plant Cell 33, 1361–1380.

126. Wijnker, E., Harashima, H., Müller, K., Parra-Nuñez, P., de Snoo, C.B., van de Belt, J., Dissmeyer, N., Bayer, M., Pradillo, M., and Schnittger, A. (2019). The Cdk1/Cdk2 homolog CDKA;1 controls the recombination landscape in Arabidopsis. Proc Natl Acad Sci U S A 116, 12534–12539.

127. Willis, K.J., Feurdean, A., Birks, H.J., Bjune, A.E., Breman, E., Broekman, R., Grytnes, J.A., New, M., Singarayer, J.S., and Rozema, J. (2011). Quantification of UV-B flux through time using UV-B-absorbing compounds contained in fossil Pinus sporopollenin. New Phytologist 192, 553–560.

128. Wong, R., and Forer, A. (2004). Backward chromosome movement in crane-fly spermatocytes after UV microbeam irradiation of the interzone and a kinetochore. Cell Biology International 28, 293–298.

129. Xiong, S.-X., Lu, J.-Y., Lou, Y., Teng, X.-D., Gu, J.-N., Zhang, C., Shi, Q.-S., Yang, Z.-N., and Zhu, J. (2016). The transcription factors MS188 and AMS form a complex to activate the expression of CYP703A2 for sporopollenin biosynthesis in Arabidopsis thaliana. Plant Journal 88, 936–946.

130. Xu, C., Liu, Z., Zhang, L., Zhao, C., Yuan, S., and Zhang, F. (2013). Organization of actin cytoskeleton during meiosis I in a wheat thermo-sensitive genic male sterile line. Protoplasma 250, 415–422.

131. Xu, Y., Charles, M.T., Luo, Z., Mimee, B., Tong, Z., Véronneau, P.Y., Roussel, D., and Rolland, D. (2019). Ultraviolet-C priming of strawberry leaves against subsequent Mycosphaerella fragariae infection involves the action of reactive oxygen species, plant hormones, and terpenes. Plant Cell Environment 42, 815–831.

132. Xue, J.-S., Zhang, B., Zhan, H., Lv, Y.-L., Jia, X.-L., Wang, T., Yang, N.-Y., Lou, Y.-X., Zhang, Z.-B., Hu, W.-J., Gui, J., Cao, J., Xu, P., Zhou, Y., Hu, J.-F., Li, L., and Yang, Z.-N. (2020). Phenylpropanoid derivatives are essential components of sporopollenin in vascular plants. Molecular Plant 13, 1644–1653.

133. Xue, S., Zang, Y., Chen, J., Shang, S., Gao, L., and Tang, X. (2022). Ultraviolet-B radiation stress triggers reactive oxygen species and regulates the antioxidant defense and photosynthesis systems of intertidal red algae Neoporphyra haitanensis. Frontiers in Marine Science 9.

134. Xue, Z., Liu, C., Shi, W., Miao, Y., Shen, Y., Tang, D., Li, Y., You, A., Xu, Y., Chong, K., and Cheng, Z. (2019). OsMTOPVIB is required for meiotic bipolar spindle assembly. Proc Natl Acad Sci U S A 116, 15967–15972.

135. Yamamoto, A., and Hiraoka, Y. (2003). Monopolar spindle attachment of sister chromatids is ensured by two distinct mechanisms at the first meiotic division in fission yeast. EMBO Journal 22, 2284–2296.

136. Yang, C., Hu, B., Portheine, S.M., Chuenban, P., and Schnittger, A. (2020). State changes of the HORMA protein ASY1 are mediated by an interplay between its closure motif and PCH2. Nucleic acids research 48, 11521–11535.

137. Yang, C., Sofroni, K., Hamamura, Y., Hu, B., Elbasi, H.T., Balboni, M., Chu, L., Stang, D., Heese, M., and Schnittger, A. (2022). ZYP1-mediated recruitment of PCH2 to the synaptonemal complex remodels the chromosome axis leading to crossover restriction. Nucleic acids research 50, 12924–12937.

138. Yang, W., Yao, D., Duan, H., Zhang, J., Cai, Y., Lan, C., Zhao, B., Mei, Y., Zheng, Y., Yang, E., Lu, X., Zhang, X., Tang, J., Yu, K., and Zhang, X. (2023). Maize and Arabidopsis VAMP726 confers pollen resistance to heat and UV radiation by influencing lignin content in sporopollenin. Plant Communications, 100682.

139. Yi, J., Kradolfer, D., Brownfield, L., Ma, Y., Piskorz, E., Köhler, C., and Jiang, H. (2023). Meiocyte size is a determining factor for unreduced gamete formation in Arabidopsis thaliana. New Phytologist 237, 1179–1187.

140. Yin, R., Skvortsova, M.Y., Loubéry, S., and Ulm, R. (2016). COP1 is required for UV-B-induced nuclear accumulation of the UVR8 photoreceptor. Proc Natl Acad Sci U S A 113, E4415–4422.

141. Yokawa, K., and Baluška, F. (2015). Pectins, ROS homeostasis and UV-B responses in plant roots. Phytochemistry 112, 80–83.

142. Yoshiyama, K.O., Kaminoyama, K., Sakamoto, T., and Kimura, S. (2017). Increased phosphorylation of Ser-Gln sites on SUPPRESSOR OF GAMMA RESPONSE1 strengthens the DNA damage response in Arabidopsis thaliana. Plant Cell 29, 3255–3268.

143. Yu, S., Galvão, V.C., Zhang, Y.-C., Horrer, D., Zhang, T.-Q., Hao, Y.-H., Feng, Y.-Q., Wang, S., Schmid, M., and Wang, J.-W. (2012). Gibberellin regulates the Arabidopsis floral transition through miR156-targeted SQUAMOSA PROMOTER BINDING–LIKE transcription factors. Plant Cell 24, 3320–3332.

144. Zeng, Q.N., Chen, J.G., and Ellis, B.E. (2011). AtMPK4 is required for male-specific meiotic cytokinesis in Arabidopsis. Plant Journal 67, 895–906.

145. Zhang, C., Yang, Y.P., and Duan, Y.W. (2014). Pollen sensitivity to ultraviolet-B (UV-B) suggests floral structure evolution in alpine plants. Scientific Reports 4, 4520.

146. Zhang, C., Shen, Y., Tang, D., Shi, W., Zhang, D., Du, G., Zhou, Y., Liang, G., Li, Y., and Cheng, Z. (2018). The zinc finger protein DCM1 is required for male meiotic cytokinesis by preserving callose in rice. Plos Genetics 14, e1007769.

147. Zhang, Q., Lin, F., Mao, T., Nie, J., Yan, M., Yuan, M., and Zhang, W. (2012). Phosphatidic acid regulates microtubule organization by interacting with MAP65-1 in response to salt stress in Arabidopsis. Plant Cell 24, 4555–4576.

148. Zhang, Q., Lin, L., Fang, F., Cui, B., Zhu, C., Luo, S., and Yin, R. (2023). Dissecting the functions of COP1 in the UVR8 pathway with a COP1 variant in Arabidopsis. Plant Journal 113, 478–492.

149. Zhang, X., Ling, Y., Wang, W., Zhang, Y., Ma, Q., Tan, P., Song, T., Wei, C., Li, P., Liu, X., Ma, R.Z., Zhong, H., Cao, C., and Xu, Q. (2013). UV-C irradiation delays mitotic progression by recruiting Mps1 to kinetochores. Cell cycle (Georgetown, Tex.) 12, 1292–1302.

150. Zhao, J., Gui, X., Ren, Z., Fu, H., Yang, C., Wang, W., Liu, Q., Zhang, M., Wang, C., Schnittger, A., and Liu, B. (2023). ATM-mediated double-strand break repair is required for meiotic genome stability at high temperature. Plant Journal 114, 403–423.

151. Zhou, Q., Cheng, X., Kong, B., Zhao, Y., Li, Z., Sang, Y., Wu, J., and Zhang, P. (2022). Heat shock-induced failure of meiosis I to meiosis II transition leads to 2n pollen formation in a woody plant. Plant Physiology 189, 2110–2127.

152. Zickler, D., and Kleckner, N. (2023). Meiosis: dances between homologs. Annu Rev Genet 57, 1–63.

